# Tissue architectural cues drive organ targeting of human tumor cells in zebrafish

**DOI:** 10.1101/233361

**Authors:** Colin D. Paul, Kevin Bishop, Alexus Devine, Elliott L. Paine, Jack R. Staunton, Sarah M. Thomas, Lisa M. Miller Jenkins, Nicole Y. Morgan, Raman Sood, Kandice Tanner

## Abstract

Sites of metastasis are non-random, with certain types of cancers showing organ preference during distal colonization. Using multiple brain- and bone marrow-seeking human and murine breast cancer subclones, we determined that tumor cells that home to specific murine organs (brain and bone marrow) ultimately colonized analogous tissues (brain and caudal vein plexus [CVP]) in larval zebrafish. We then exploited the zebrafish model to delineate factors leading to differential cell homing and extravasation. Bone marrow-tropic clones showed higher expression of integrins and focal adhesions associated with mechanosensing machinery than brain-tropic clones and were more sensitive to vessel topography during extravasation. Knockdown of β1 integrin reduced extravasation and redistributed organ targeting from disordered vessels in the CVP to the brain. Our results show that organ selectivity is driven by topography- and cell type-dependent extravasation at the tumor-endothelial interface in the larval zebrafish and provide important insights into the early stages of metastasis.

## INTRODUCTION

Metastasis describes the process whereby cancer cells move from a primary tumor and establish lesions in distinct organs (Nguyen et al., 2009; Obenauf and Massague, 2015; Steeg, 2016). Metastasis is non-random, with differences in organ targeting and clinical latency correlated with initial tumor type in line with Paget’s seed and soil hypothesis (Nguyen et al., 2009; Obenauf and Massague, 2015; Paget, 1989; Steeg, 2016). Differences in tumor targeting and latency following dissemination are critical for the treatment of certain types of cancer (Kennecke et al., 2010; Steeg, 2016; Steeg et al., 2011; Tanner and Gottesman, 2015). For example, breast cancer cells can disseminate through the circulation prior to the primary tumor reaching a clinically detectable size, and latent disseminated cells can lead to re-emergence of aggressive cancers up to ~20 years following initial treatment (Steeg et al., 2011). Therefore, understanding how cells target specific organs, whether differences exist in this targeting, and the factors critical to cell survival following dissemination are critical for developing treatments for metastatic and refractory cancers.

A limited repertoire of isogenic clones exists such that multiple organ-seeking clones are derived from the same parental clone within the same strain of mouse (Nguyen et al., 2009; Yoneda et al., 2001; Zhang et al., 2013). From these model systems, cell (“seed”) intrinsic differences in genes and signaling pathways that regulate organ specificity have been evaluated (Nguyen et al., 2009; Yoneda et al., 2001; Zhang et al., 2013). Additionally, understanding of the mechanisms of tumor cell dissemination from the primary site and evaluation of the abundance of circulation tumor cells in the bloodstream have made it clear that cell entry into the circulatory system is vital for metastatic outgrowth (Condeelis and Segall, 2003; Ewald et al., 2011; Friedl and Alexander, 2011; Hirata et al., 2015). However, the mechanisms that determine how cells transition from the circulation to successfully colonize the “soil” at distant organs is less understood, particularly in the context of the earliest stages of metastasis and simultaneously in multiple organs.

An important mediator of tumor cell fate that interacts with cell-intrinsic genetics is the microenvironment (Bissell and Hines, 2011; Friedl and Alexander, 2011; Tanner and Gottesman, 2015). Tumor cells along the metastatic cascade encounter different physical microenvironments during capillary arrest and extravasation into organs (Kim and Tanner, 2015; Kumar and Weaver, 2009). Specifically, as cells traffic through conduits in the lymphatic and blood circulation system, both the size and curvature of the blood vessels can vary. Numerous in vitro studies have demonstrated how topographical cues can influence cell motility, invasion, and gene expression (Paul et al., 2017). We thus hypothesized that differences in the physical microenvironment in distinct tissues as cells transition from the bloodstream to the organ parenchyma is a key determinant of organ specificity. These steps in early colonization are rare and difficult to study using current models (Azevedo et al., 2015). However, the relatively small size and optical transparency of the larval zebrafish enables imaging of multiple tissues in the same animal at single cell resolution. The zebrafish is rapidly becoming a model for studying tumor behavior at different stages of the metastatic cascade (Stoletov et al., 2010; Stoletov et al., 2007; White et al., 2013), and several organs important for human metastasis are sufficiently conserved across vertebrates. These organs share similar cell types, tissue architecture, immune regulation, and ECM composition with mammalian organs. Within these organ environments, the vascular network is comprised of a myriad of branching and anastomosing blood and lymphatic vessels. Thus, using this system, we can also interrogate the role of vessel sizes and architecture in early colonization and evaluate how vessel topographical cues may drive organ targeting and extravasation.

Here, we determined that brain- or bone/bone marrow-targeting human and murine breast tumor cells showed non-random organ targeting following direct injection into the zebrafish circulation. Specifically, cell lines that preferentially targeted bone marrow and brain niches in mice also show similar targeting in the larval fish. Differences in extravasation emerged at organs of different topographic complexity for different cell lines, indicating that topography was a key cue in organ targeting. Silencing of β1 integrin redirected cell homing and differentially impacted extravasation in tissues of varying vessel topography. Overall, these data suggest that patterns of metastatic spread are driven, at least in part, by topography-driven patterns of extravasation during early metastatic dissemination.

## RESULTS

### Human cancer cells successfully undergo early metastatic colonization within multiple tissues of the zebrafish

The metastatic cascade involves cell invasion or shedding into the circulatory system, arrest in the capillaries of target organs, extravasation into the parenchyma of these organs, and either dormancy or proliferation to establish a secondary lesion (Kim and Tanner, 2015; Kumar and Weaver, 2009). Therefore, we first asked whether human cancer cells could complete these steps to colonize multiple tissues within the zebrafish following transit through the circulatory system. We injected human cancer cells directly into the circulation of 2 days post-fertilization (dpf) zebrafish, which were imaged at 5 days post-injection (dpi; zebrafish age of 7 dpf) to assess organ colonization (Figure 1A).

**Figure 1.**
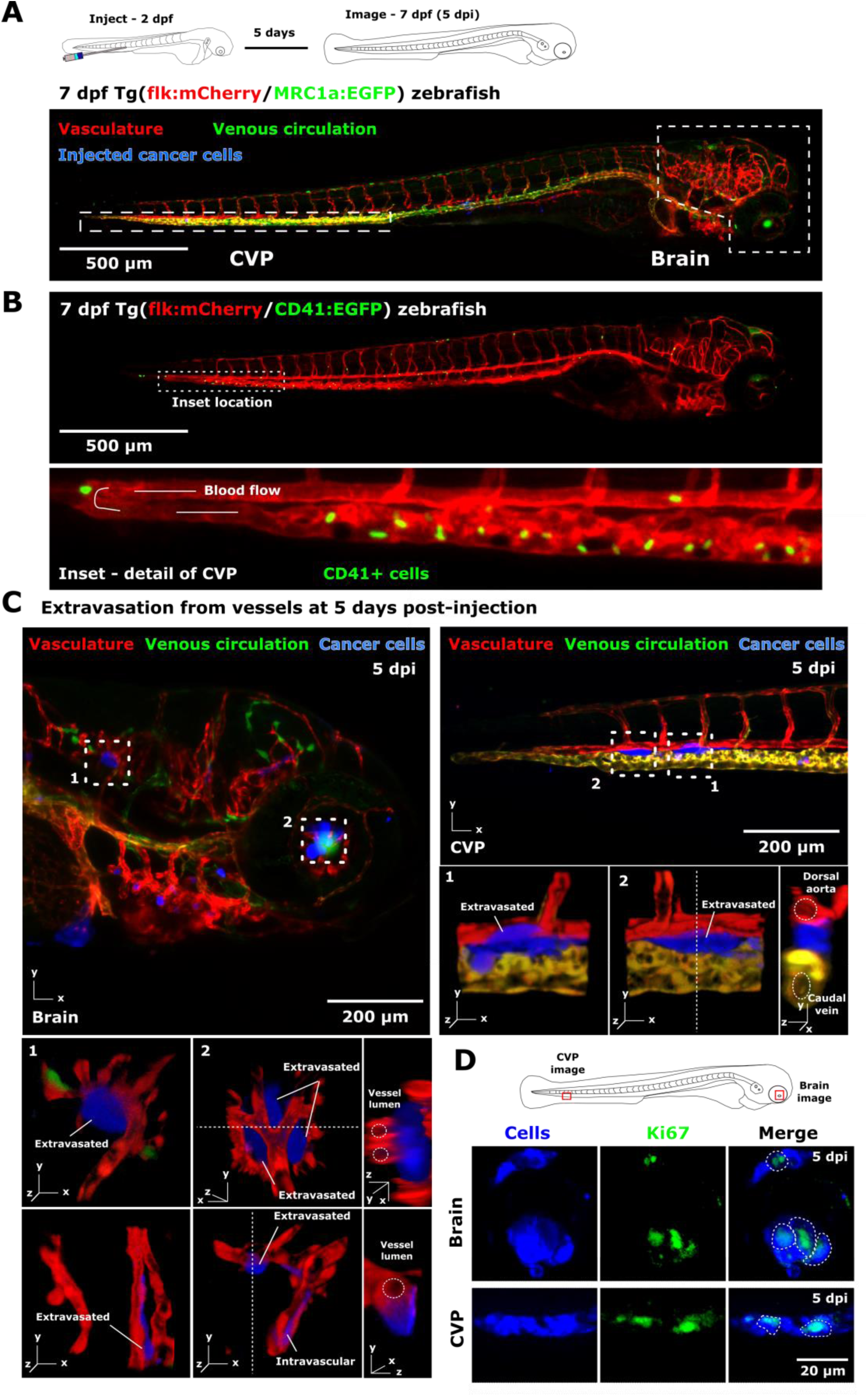
Injected cancer cells recapitulate key steps of the early metastatic cascade in a zebrafish xenograft model. (A) Schematic of experimental protocol. Cancer cells (MDA-MB-231 brain-tropic [231BR]) were injected into the circulation of transgenic Tg(flk:mCherry/MRC1a:EGF) zebrafish at 2 days post-fertilization (dpf) via the dorsal aorta. Fish were imaged at 5 days post-injection (dpi) at an age of 7 dpf. An overview (tiled average intensity projection from confocal z stack) of a 7 dpf zebrafish is shown for context. The vasculature is displayed in red, the venous circulation in green, and injected cancer cells in blue. The brain and caudal vein plexus (CVP) are outlined. Scale is indicated. (B) Transgenic Tg(CD41:EGFP/flk:mCherry) zebrafish with fluorescently labeled CD41-positive cells and vasculature at 7 dpf. Image displays average intensity projections from confocal z stack. Inset shows detail of the sinusoidal vessel architecture and CD41-positive population in the CVP. The vasculature is displayed in red, and CD41-positive cells in green. Arrows indicate direction of blood flow. (C) Detail of cancer cells in the zebrafish brain (left panels, 231BR cells) and CVP (right panels, MDA-MB-231 bone marrow-tropic [231BO] cells) at 5 dpi. Regions of interest (labeled 1,2) are indicated on overview images of the brain and CVP (average intensity projections from confocal z stacks). 3D reconstructions of these regions are shown in detail below. Two additional images of cells in the brain of a different zebrafish larva are also shown. For select images, orthogonal views of extravasated cells are presented, with the vessel lumen of neighboring vessels outlined. (D) Cancer cells were injected into the circulatory system of Tg(flk:mCherry/mpx:EGFP) zebrafish at 2 dpf and fixed at 5 dpi. Cells were stained for Ki67 using whole-mount immunofluorescence, and zebrafish were imaged in the regions indicated on the schematic using confocal microscopy. Images in the brain and CVP were taken from separate larvae. Images are average intensity projections of confocal z stacks from indicated regions of interest. Individual channels and merged images of cells in the zebrafish brain (231BR cells) and CVP (231BO cells) are displayed, with cells in blue and Ki67 in green. Outlines indicate cells staining positive for Ki67. Scale is indicated. See also Supplementary Videos 1,2.

Organs that are frequent sites of metastasis in human patients are well conserved between mammals and zebrafish (Meeker and Trede, 2008; Oosterhof et al., 2015). Brain and bone marrow niches are common sites of metastasis in patients presenting with breast cancer. Hence, we focused on two regions of the larval zebrafish, the caudal vein plexus (CVP) and brain, that can serve as organ analogs for their mammalian counterparts at this developmental stage (Figure 1A). At the cellular level, the zebrafish larval brain shares similarities with the mammalian brain in that the cellular components, for example, microglia, astrocytes, neurons, and blood vessels (Oosterhof et al., 2015), are conserved, as well as key architectural features, such as a functional blood-brain barrier (Fleming et al., 2013) and choroid vessel. The CVP is comprised of sinusoidal vessels which surround the caudal hematopoietic tissue (CHT), which is analogous to mammalian fetal liver or bone marrow and harbors hematopoietic stem cells (HSCs), including CD41 positive cells (Figure 1B). We observed, in both the brain and CVP, a population of cells that were able to extravasate and survive outside of vessels in both tissues at 5 dpi (Figure 1C; Supplementary Videos 1,2). Extravasated cells often maintained a perivascular position and could be observed both as single cells or as groups of cells (Figure 1C; Supplementary Videos 1,2). A subset of cells was active in the cell cycle by Ki67 staining (Figure 1D), indicating that this model successfully recapitulated extravasation and early colonization for multiple organs. Confident that cells were able to successfully colonize multiple tissues of the larval zebrafish, we next took advantage of the model to study early steps in organ targeting.

### Organotropic human tumor cells show non-random distal organ colonization in a zebrafish model of metastasis

As the key components of the zebrafish brain and hematopoietic niche have emerged and are functional in larval zebrafish in the first few days after fertilization (Fleming et al., 2013; Goessling and North, 2011; Murayama et al., 2006; Oosterhof et al., 2015), we reasoned that at this early developmental stage, there might be sufficient conservation between mammalian and zebrafish tissue to examine organ targeting in the larval zebrafish. We hypothesized that human tumor cells that home to murine organs (Kim et al., 2018; Kusuma et al., 2012; Yoneda et al., 2001; Zhang et al., 2013) may show similar patterns of dissemination and survival in larval zebrafish. Sub-clones of breast cancer cells that were selected for their potential to home to murine brain and bone were injected into the circulatory system of 2 dpf zebrafish via injection to the dorsal aorta (Figure 2A). We quantified cell homing to the brain and CVP after a targeting window of 1-5 days (Figure 2A). Cell homing encompassed quantification of all cells present (i.e., both intravascular and extravascular cells) in a particular organ for each of the pairs of clones and was assessed on a per larva basis, as we were able to image both the brain and CVP of individual animals with this model system (Figure 2A). This enabled cell-scale analysis of differential colonization patterns during the earliest stages of metastasis.

**Figure 2.**
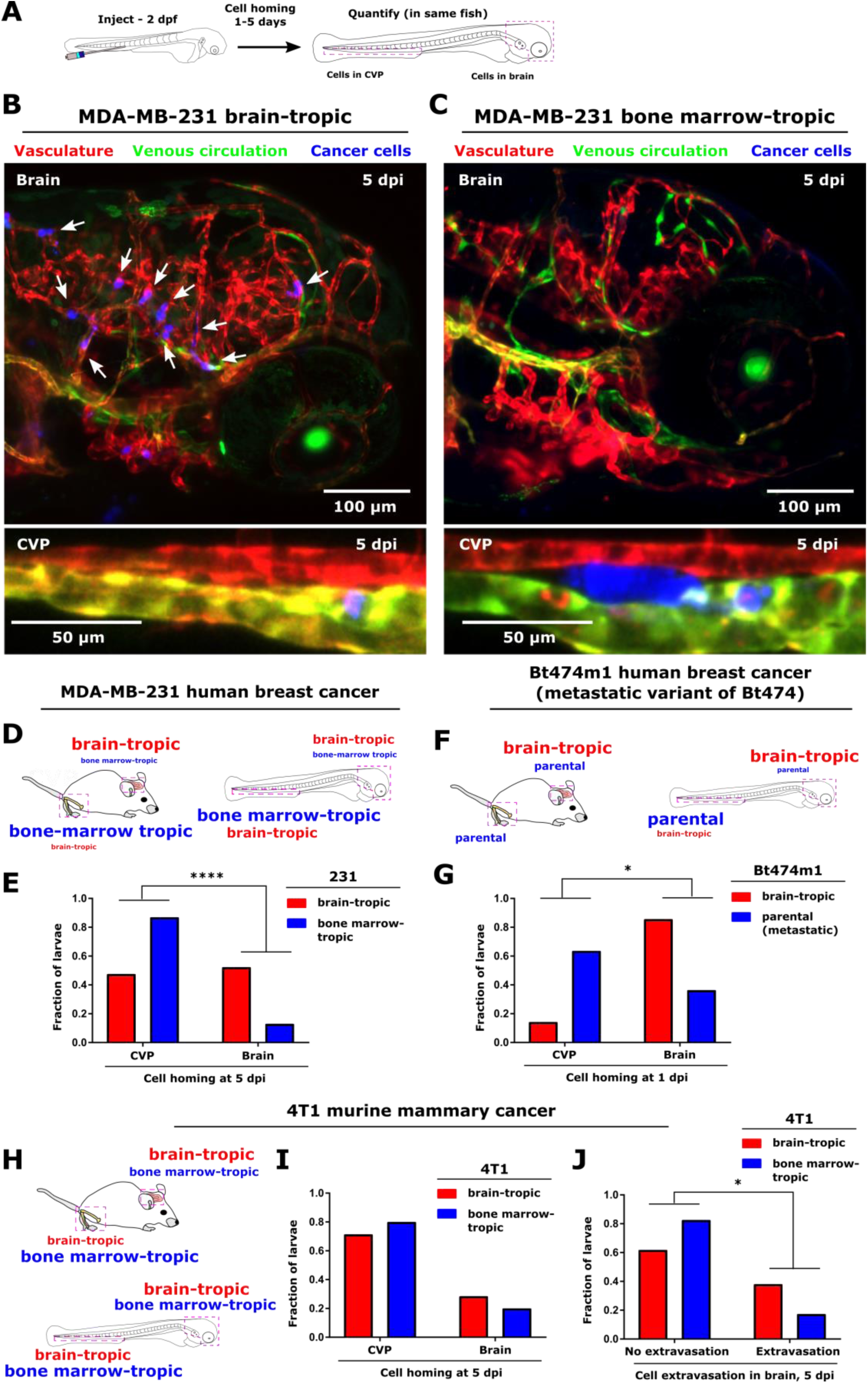
Non-random targeting of organ-tropic human cell lines in a zebrafish metastasis model. (A) Schematic of experimental setup. Cancer cells were injected into the circulatory system of Tg(flk:mCherry/MRC1a:EGFP) zebrafish at 2 days post-fertilization (dpf) and imaged at 1–5 days post-injection (dpi; zebrafish age at imaging was 7 dpf). Cell homing was assessed by quantifying the relative number of cells present in the brain and caudal vein plexus (CVP) region of each larva. Representative images of MDA-MB-231 (C) brain-tropic subclone (231BR) and (D) bone marrow-tropic subclone (231BO) colonization in the zebrafish brain and CVP at 5 dpi. Arrows indicate cell positions in the brain. Images are average intensity projections from confocal z stacks. The vasculature is displayed in red, the venous circulation in green, and injected cancer cells in blue. Scales are indicated. (D) Schematic of reported organ colonization of MDA-MB-231 brain-tropic (231BR, red) and bone marrow-tropic (231BO, blue) subclones in murine models, and of the observed cell homing in the zebrafish model. Font size indicates relative homing within a given organ. (E) Distribution of zebrafish larvae demonstrating homing to the CVP and brain at 5 dpi for MDA-MB-231 organ-targeting clones. ****, p<0.0001 by two-tailed binomial distribution of brain-tropic cells compared to the observed distribution of bone-marrow tropic cells. N=21 zebrafish injected with 231 brain-tropic cells (398 total cells), N=23 zebrafish injected with 231 bone marrow-tropic cells (376 total cells). (F) Schematic of reported organ colonization of Bt474m1 cells (blue) and brain-tropic subclones (Bt474m1-Br3, red) in murine models, and of the observed cell homing in the zebrafish model. Font size indicates relative homing within a given organ. (G) Distribution of zebrafish larvae demonstrating homing to the CVP and brain at 1 dpi for Bt474m1 clones. *, p<0.05 by two-tailed binomial distribution of brain-tropic cells compared to the observed distribution of parental cells. N=7 zebrafish injected with brain-tropic cells (53 total cells), N=11 zebrafish injected with parental Bt474m1 cells (64 total cells). (H) Schematic of reported organ colonization of 4T1 brain-tropic (4T1Br4, red) and bone marrow-tropic (4T1BM2, blue) subclones in murine models, and of the observed cell homing in the zebrafish model. Font size indicates relative homing within a given organ. (I) Distribution of zebrafish larvae demonstrating homing to the CVP and brain at 5 dpi for 4T1 organ-targeting clones. Distribution of larvae between cell types was not statistically significant. N=28 zebrafish injected with brain-tropic cells (316 total cells), N=25 zebrafish injected with bone marrow-tropic cells (153 total cells). (J) Fraction of larvae from the population examined in panel I demonstrating extravasation in the brain at 5 dpi for 4T1 organ-targeting clones. N=21 zebrafish for brain-tropic cell injections, N=23 zebrafish for bone marrow-tropic cell injections. Cells with one cell extravasated in the brain at 5 dpi were excluded from analysis. *, p<0.05 by two-tailed binomial test, with the expected fraction of larvae exhibiting no extravasation or extravasation set by the bone marrow-tropic line. See also Figures S1-S3, Supplementary Video 3.

Specifically, cell homing of brain-targeting cells to the brain was quantified for animals injected with MDA-MB-231 human breast adenocarcinoma cells that homed to murine brain vs. bone marrow (Yoneda et al., 2001), and a significantly higher fraction of larvae demonstrated homing to the brain when injected with brain-tropic 231 cells compared to bone marrow-tropic cells at 5 dpi (Figure 2B-E) Similarly, animals injected with brain-targeting sub-clones of HER2^+^ Bt474m1 human breast cancer cells (Zhang et al., 2013) were enriched for the population in which brain targeting was observed at 1 dpi compared to the cohort injected with parental Bt474m1 (a metastatic variant of Bt474) cells (Figure 2F,G; Figure S1B,C). While a fraction of larvae injected with brain-targeting MDA-MB-231 and Bt474m1 cells exhibited targeting to the CVP, we consistently observed a statistically significant increase in the fraction of larvae with cells homing to the brain compared to larvae injected with bone-targeting or parental cells (Figure 2E,G), and cell homing results observed in the zebrafish showed similar patterns to organ colonization of these cell lines observed in murine models (Figure 2D,F).

In addition, we tested an isogenic pair of brain- and bone marrow-tropic murine breast cancer cell lines. In this system, both the brain- and bone marrow-tropic clones also show a weak tropism to the bone marrow and brain, respectively, in murine models (represented schematically in Figure 2H). Quantitation of cell distributions in the zebrafish revealed that these murine breast cancer 4T1 cells home to both the brain and CVP in zebrafish (Figure 2H, I; Figure S1D,E). However, many of the bone marrow-tropic cells that homed to the brain remained in the luminal spaces of the vessels. Further quantitation of cell extravasation revealed that the brain-targeting 4T1 cells exhibited extravasation in the brain for a significantly higher fraction of injected animals (Figure 2J).

Having observed distinct organ targeting, we next asked if these observed differences were due simply to differences in survival in the tissue parenchyma at a particular metastatic site. Thus, we co-injected both MDA-MB-231 clones directly into the brain of optically transparent Casper zebrafish (Figure S2A). We determined via both sampling of fish from the injected cohort at different time points (Figure S2A, B) and longitudinal imaging of the same fish (Figure S2A, C-D) that both clones showed similar survival for one week within the brain parenchyma. To confirm that survival of the bone-tropic clone was not due to cytokines or the physical presence of the brain-tropic clone, we also performed injection of a single clone into the brain, where we observed that both brain- and bone marrow-tropic cells exhibited robust and similar cell survival when alone in the brain (Figure S2F-H).

Organ selectivity could also arise due to organ- and cell-specific differences in clearance by innate immune cells at early stages of tumor cell colonization. To test whether the innate immune response drove organ selectivity in this system, we injected MDA-MB-231 brain- and bone marrow-tropic cells into the circulation of transgenic zebrafish with fluorescently labeled neutrophils and vasculature. We observed minimal interaction of neutrophils with either clone in the brain and CVP over the first ~14 h post-injection (Supplementary Video 3). Additionally, at 1 dpi, similar numbers of zebrafish neutrophils were present in the head and tail of larvae injected with either cell type, and the distribution of neutrophils between the head and tail was not cell-type dependent (Figure S3). We therefore concluded that organ selectivity was not dependent on neutrophil-dependent cell survival.

### Differential extravasation patterns emerge at the tumor cell – endothelial interface in zebrafish in a cell type- and organ-specific manner

Having observed non-random targeting of human cancer cell lines in the larval zebrafish, we focused on MDA-MB-231 brain-tropic (231BR) and bone marrow-tropic (231BO) cells to examine factors driving this targeting. Gene signatures have been observed to be drivers of organotropism (Obenauf and Massague, 2015). To test what cell-intrinsic factors may regulate non-random targeting in the larval zebrafish, we performed label-free mass spectrometry on each cell line. The differentially regulated proteins between the cell lines included integrins, focal adhesions, and proteins associated with contractility (Figure 3A). These proteins are implicated in a wide range of cellular processes such as migration, invasion, proliferation, and mechanosensing. Microenvironmental cues such as tumor-stromal crosstalk are important regulators of disseminated tumor outgrowth (Barnes et al., 2017; Bissell and Hines, 2011; Tanner and Gottesman, 2015). From the proteomic analysis, several proteins that are important for the cells’ ability to sense physical cues from the microenvironment are differentially regulated between the brain-tropic and bone marrow-tropic clones. Hence, differential cellular response to physical cues may influence organ selectivity. Therefore, we first assessed whether differences in cell trafficking through the vessel conduits available for trafficking and migration upon cell injection influenced organ homing. Vessel diameters in the brain and intersegmental vessels (ISVs; nearly linear vessels in the zebrafish tail and trunk) averaged 9.9±2.4 and 10.6±2.6 µm, respectively, and ranged from ~5-15 µm in width (Figure 3B). These values are similar to those measured in human lung and nailbed capillaries and are comparable to those seen in mice, rats, and rabbits (Doerschuk et al., 1993; Kienast et al., 2010; Mathura et al., 2001; Potter and Groom, 1983; Tufto and Rofstad, 1999). Vessel diameters in the were CVP was more widely varied, and vessels ranged from ~5-50 µm in width with an average of 16.7±6.8 µm (Figure 3B).

**Figure 3.**
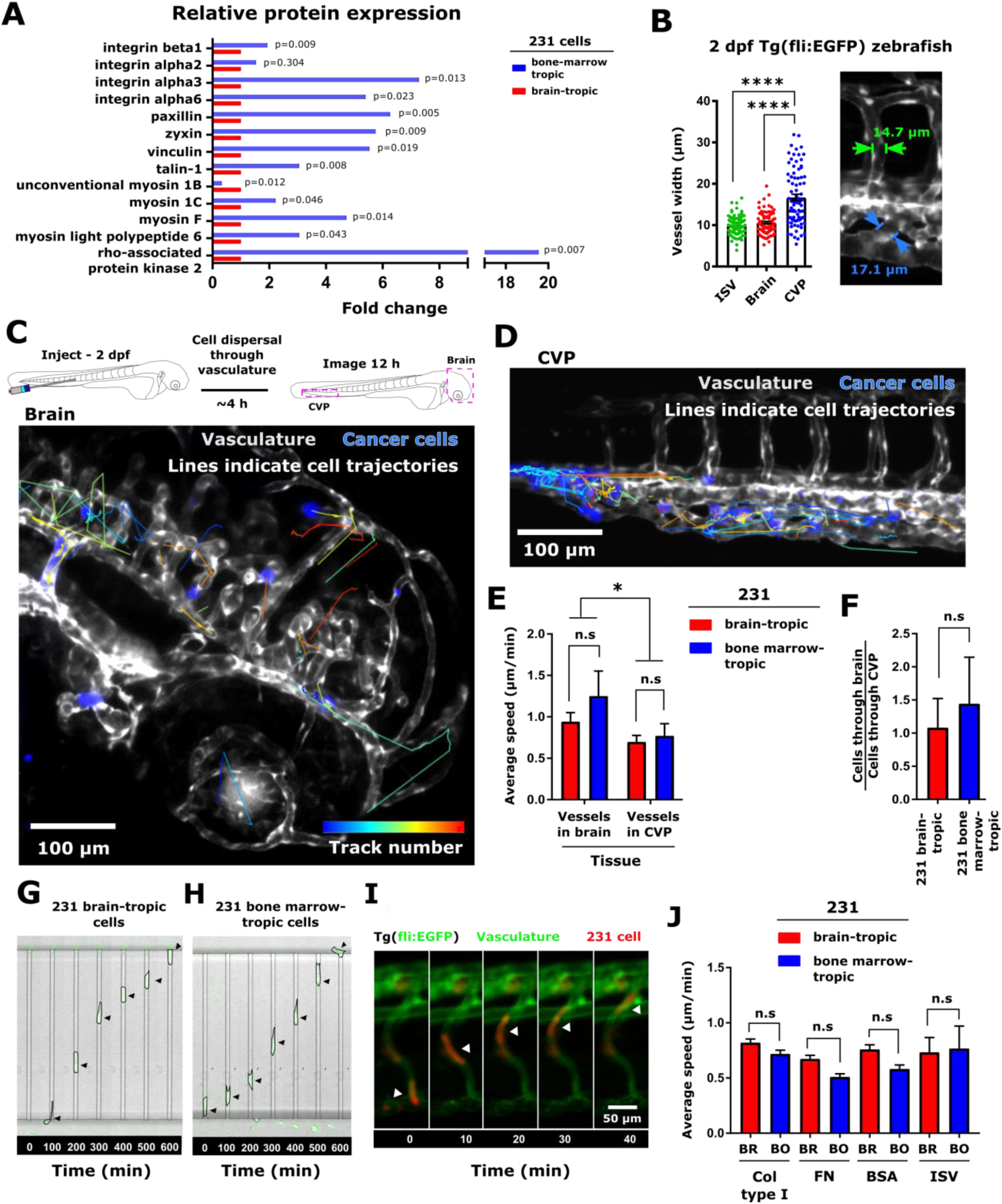
Cell speed in zebrafish organs is not differentially regulated between organ-tropic cell lines. (A) Relative protein expression of key adhesion, contractility, and mechanosensitive proteins in MDA-MB-231 organ-tropic cell lines. For each protein, expression in bone marrow-tropic cells is normalized to that in brain-tropic cells, and p-values are indicated. (B) Average ± SEM of blood vessel widths in the zebrafish intersegmental vessels (ISV), caudal vein plexus (CVP), and brain. Vessels were measured from four Tg(fli:EGFP) zebrafish at 2 dpf, 20 vessels per region/fish, to obtain a total of N=80 vessels at each location. ****, p<0.0001 by Dunn’s multiple comparisons post-test following Kruskal-Wallis test. (C) Schematic and representative results of in vivo cell tracking experiment. 2 dpf Tg(flk:mCherry/MRC1a:EGFP) zebrafish were injected with MDA-MB-231 brain-tropic (231BR) or bone marrow-tropic (231BO) cells in the dorsal aorta. The brain and (D) CVP were then imaged for 12 h post-injection to visualize cell trafficking within multiple tissues of the same fish. Representative images of cell trajectories within the zebrafish brain and CVP are shown. Trajectories are overlaid on the 12 h image. Tracks are pseudocolored by track number (blue=first track, red=final track). Cells are displayed in blue. Zebrafish vasculature and venous circulation are shown in grayscale to facilitate visualization of cell trajectories. Scale bar = 100 μm. (E) Average ± SEM speeds of cells trafficking through the brain and CVP. Speeds are grouped measurements (brain-tropic cells – 91 cells in brain, 131 cells in CVP; bone marrow-tropic cells – 46 cells in brain, 56 cells in CVP) taken from multiple fish (brain-tropic cells – 6 larvae; bone marrow-tropic cells – 8 larvae). *, p<0.05 by two-way ANOVA, with tissue but not cell type as a significant source of variation; n.s., not significant by Sidak’s multiple comparisons post-test between cell lines in each tissue. (F) Per larva ratio of cells tracked in the brain to cells tracked in the CVP for both brain-tropic and bone marrow-tropic cells. Plot displays mean ± SEM for this ratio averaged across N=7 larvae for brain-tropic cells (310 cells total) and N=10 larvae for bone marrow-tropic cells (97 cells total). Time-lapse images of MDA-MB-231 (G) brain-tropic cells and (H) bone marrow-tropic cells migrating through confining microchannels coated with 20 μg/ml rat tail collagen type I. Arrows indicate cell position. For scale, microchannel width = 8 µm. (I) Representative image of MDA-MB-231 cell (red) migrating through Tg(fli:EGFP) zebrafish ISV (green). Arrow indicates cell location. Scale is indicated. (J) Average speeds (mean ± SEM) of cells migrating in microchannels and zebrafish intersegmental vessels. In the microchannels, speeds from at least 52 cells collected over at least 2 biological duplicates were measured. Speeds in the intersegmental vessels were measured for N=38 brain-tropic cells from 6 larvae and N=14 bone marrow-tropic cells from 7 larvae. For a given microchannel protein coating and within the ISV, speeds were not significantly different (n.s.) by Sidak’s multiple comparisons post-test following two-way ANOVA. See also Figure S4 and Supplementary Video 4.

We then assessed cell motility in vivo in the brain and CVP. Intravital imaging began from ~4-6 h after injection and continued for the subsequent 12 h (Figure 3C). Tracking of cell movement in 3D through vessels in the zebrafish brain and CVP revealed that cells moved at greater average speeds within the lumen of vessels in the brain than within the CVP for both clones, where the organ of observation was a statistically significant source of variation in cell speed by two-way ANOVA (Figures 3C-E; Supplementary Video 4). However, speeds did not differ significantly between the cell types within a given organ (Figure 3E). Observed speeds were several orders of magnitude less than the blood velocity in larval zebrafish (Watkins et al., 2012) and were not due to zebrafish growth, which was corrected for using 3D image registration (Parslow et al., 2014) (Figure S4). On average, an equal number of cells passed through the brain and CVP (normalized per fish, where both organs could be imaged), indicating that any observed differences in cell speed or occlusion were not driven by unequal exposure to cells during their movement through the circulation (Figure 3F).

One concern may be that the signaling cues that drive cross talk between host and tumor cells may not be fully conserved in the human-zebrafish xenograft system, and that the similar speeds observed for the subclones were therefore an artifact of the experimental system. Thus, we fabricated in vitro microchannel mimetics that recreated vessel widths but where the luminal spaces were coated with extracellular matrix (ECM) proteins. Comparison of cell migration within the geometric in vitro mimetics (Figure 3G, H) and in vivo within zebrafish ISVs (Figure 3I) revealed that both clones showed similar motilities during confined migration (Figure 3J).

During transit, cell arrest, and extravasation, cells receive cues from the architecture of the surrounding blood and lymph vessels in addition to confinement due to vessel width and tissue mechanics. At the single cell level, a cell senses its environment using protrusions that may span a few microns (filopodia) to roughly the diameter of a cell (blebs and lamellipodia). Thus, to determine the topography a cell would sense in each microenvironment, we calculated an order parameter describing the tortuosity of the vessel on the length scale corresponding to roughly that of a cell diameter (20 µm) (Figure 4A) (Fonck et al., 2009). The order parameter was measured at 2 dpf, the age at which cancer cells were introduced to the fish (Figure 4B). The order parameter in the highly oriented intersegmental vessels dorsal to the CVP was close to 1, while the order parameter was significantly lower in the CVP compared to both the brain and ISVs. Thus, the ISVs were the most ordered, whereas the CVP vessels had the greatest tortuosity and the brain vessels showed intermediate order (Figure 4B-D).

**Figure 4.**
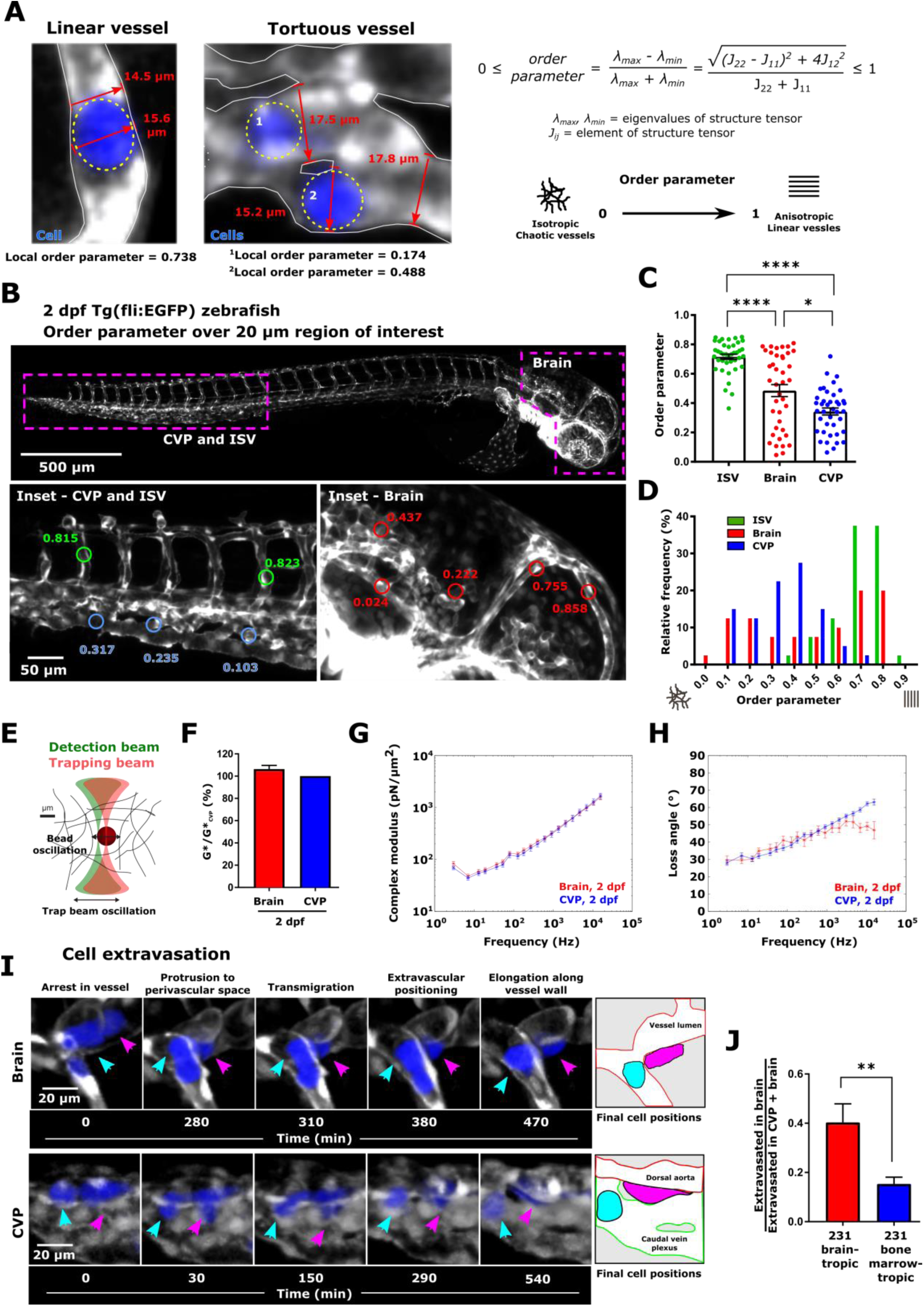
Topographical determination of extravasation at the tumor-endothelial cell interface. (A) Representative images of linear and tortuous zebrafish blood vessels. Cancer cells are shown in blue and outlined by dashed yellow ovals. Representative examples of cell and vessel width measurements are illustrated. Additionally, the local order parameter describing vessel topography as defined is given in the vicinity of each indicated cell. (B) Overview images and representative order parameter measurements in the intersegmental vessels (ISV), caudal vein plexus (CVP), and brain of 2 days post-fertilization (dpf) Tg(fli:EGFP) zebrafish. Order parameters were measured over 20 µm-diameter circular regions of interest. (C) Average ± SEM and (D) distribution of the order parameter characterizing the topography of vessels in the ISV, brain, and CVP over 20 µm-diameter circular regions of interest. A value of 1 indicates aligned structures, while a value of 0 indicates isotropic structures. Vessels were measured from four Tg(fli:EGFP) zebrafish at 2 dpf, 10 regions per tissue/fish, to obtain a total of N=40 vessels at each location. *, p<0.05; ****, p<0.0001 by Dunn’s multiple comparisons post-test following one-way ANOVA. (F) Schematic of optical trap-based microrheology setup. (G) Mean complex modulus in the brain and CVP at 2 dpf, normalized to the mean complex modulus in the CVP on the same day. *, p<0.05 by Tukey’s honestly significant difference test following two-way ANOVA. (H) Complex modulus (mean ± standard deviation) vs. frequency curves obtained using optical trap-based microrheology in zebrafish brain and CVP at 2 dpf. (I) Loss angle (mean ± standard deviation) vs. frequency curves obtained using optical trap-based microrheology in the zebrafish brain and CVP at 2 dpf. For panels (G-I), Samples were measured in triplicate with at least 30 beads per sample measured. (J) Time-lapse images of cells extravasating in the zebrafish brain (MDA-MB-231 brain-tropic cell) and CVP (MDA-MB-231 bone marrow-tropic cell). Magenta arrows indicate extravasating cells. Cyan arrows indicate cells that remain intravascular. Schematic illustrates final positions of cells. (K) Average ± SEM of extravasated cells in the brain divided by total number of extravasated cells in the brain and CVP for MDA-MB-231 brain- and bone marrow-tropic cells. Values were calculated on a per larva basis for N=21 larvae injected with brain-tropic cells (from 398 total cells) and N=22 larvae injected with bone-tropic cells (from 373 total cells). **, p<0.01 by Welch’s t test. See also and Supplementary Video 5.

Functionally, the physical microenvironment around cells is also characterized by microenvironmental mechanics. To test whether stiffness would contribute to the observed cell homing, we employed in *vivo* optical-trap-based active microrheology (Figure 4E) to measure tissue mechanics (Blehm et al., 2016; Staunton et al., 2017; Staunton et al., 2016a). The stiffness and viscoelasticity of the zebrafish brain and CVP are nearly identical over a wide range of frequencies at the time of cell injection at 2 dpf (approximately 50-1000 Pa; Figure 4F-H), indicating that mechanical differences did not drive differences in initial cell homing. Of note, the zebrafish tissue is similar in mechanical properties to the mammalian brain, mouse mammary fat pad, and hydrogels commonly used for 3D culture systems, such as Matrigel and HA gels mimicking basement membrane and brain ECM microenvironments, respectively (Blehm et al., 2016; Kim et al., 2016; Staunton et al., 2017; Staunton et al., 2016a).

Having observed organ-specific topographical but not micromechanical differences and differences in the cellular machinery associated with mechanosensing, we quantified whether there were organ-dependent differences in extravasation between clones. Extravasation is, by nature, an event that takes place at the endothelial barrier, with arrested cells protruding into the perivascular space, transmigrating through vessels, and establishing themselves in the tissue parenchyma (Figure 4I; Supplementary Video 5). Indeed, this was the case, as brain-tropic cells exhibited significantly greater extravasation in the brain relative to their extravasation in the CVP, while bone marrow-tropic cells were more likely to extravasate in the CVP (Figure 4J).

### Extravasation is mediated by β1 integrin in the CVP

β1 integrin has been implicated as one of the key proteins for cell fate decisions on time scales spanning hours-days when cells encounter a new environment (Tanner et al., 2012; Weaver et al., 1997). Of the canonical signaling pathways differentially regulated between brain and bone marrow tropic cells from the mass spectrometry data, β1 integrin emerged as a key integration point, with 7 of the top 10 differentially regulated canonical signaling pathways involving this integrin (Figure 5A). Therefore, we silenced β1 integrin to test the effect on cell survival and extravasation. Cells were transfected 2 days prior to injection to the zebrafish or cell lysate collection for analysis by mass spectrometry (Figure 5B). Mass spectrometry analysis revealed that β1 integrin was significantly lowered by knockdown (Figure 5B). Examination of homing to specific organs revealed that of siβ1 integrin MDA-MB-231 brain- and bone marrow-tropic cells were redirected cells toward the brain compared to cells transfected with a non-targeting control siRNA (Figure 5C, D; Figure S5A). We also observed a decrease in overall cell survival for both clones upon knockdown (Figure S5B). Decreased survival may be due in part to deficiencies in extravasation. Thus, we tested if the cells could extravasate into each organ. In the CVP, the fraction of cells that extravasated in the CVP upon knockdown was significantly reduced compared to control cells for both clones (Figure 5E). We therefore reasoned that β1 integrin function was crucial for cell extravasation and survival in the CVP for both cell types tested. In contrast, MDA-MB-231 brain-tropic cells were still able to extravasate in the brain following knockdown, albeit to a lesser extent than control cells (Figure 5F). This phenomenon was distinct from that in bone marrow-tropic cells, where knockdown of β1 integrin again eliminated extravasation in the brain (Figure 5F).

**Figure 5.**
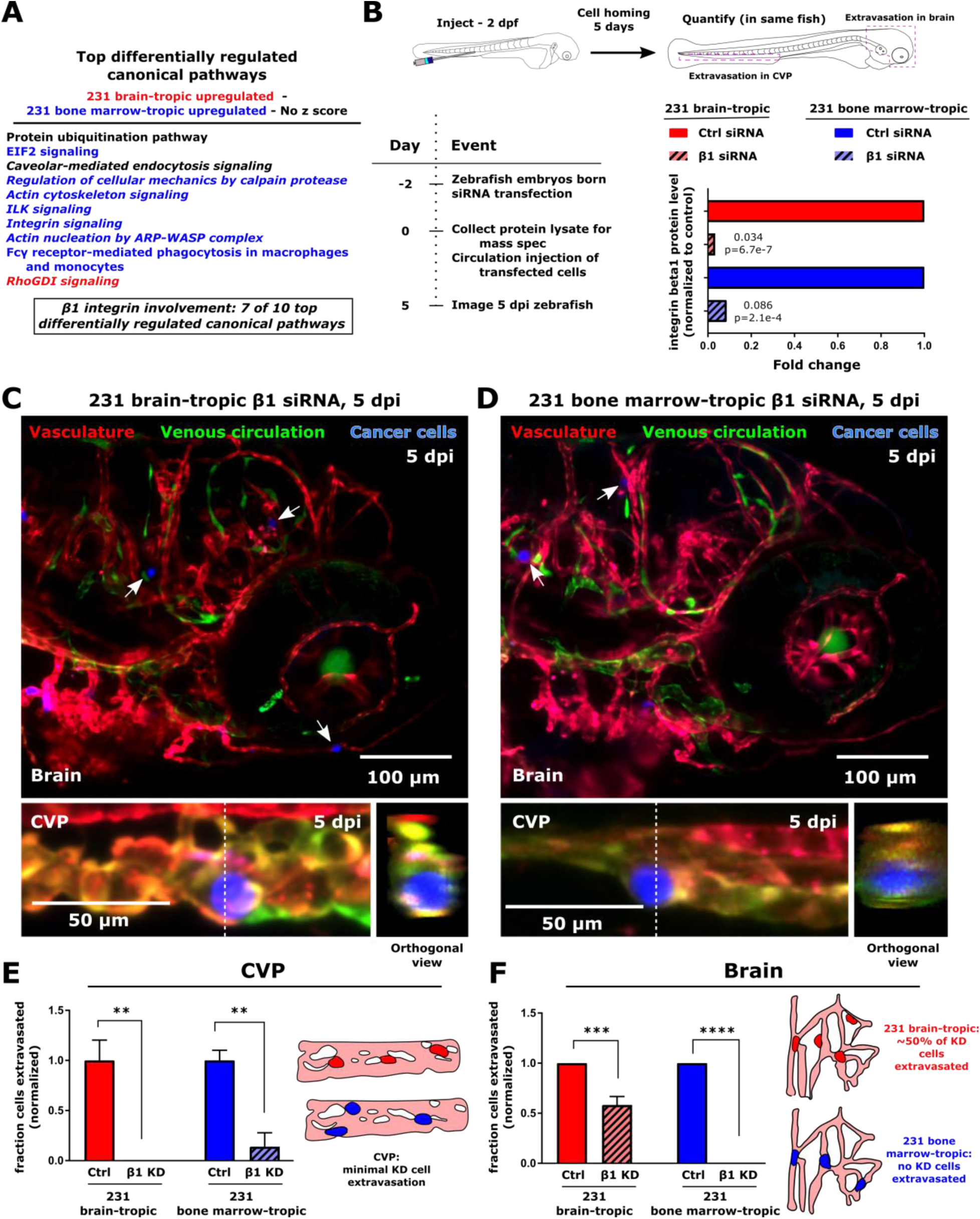
β1 integrin mediates extravasation in an organ- and cell type-specific manner. (A) List of the top differentially regulated canonical signaling pathways between MDA-MB-231 brain-tropic (231BR) and bone marrow-tropic (231BO) cells from pathway analysis of mass spectrometry data. Where possible, the direction of pathway regulation (from IPA analysis z score) is indicated (red – upregulated in 231BR; blue – upregulated in 231BO; black – no conclusive direction of regulation). Pathways in which β1 integrin are implicated are italicized. (B) Schematic and outline of siRNA knockdown experiments, and mass spectrometry results showing extent of knockdown at the time of cell injection. Cells were injected at 2 days post-fertilization (dpf), and extravasation in the brain and caudal vein plexus (CVP) was quantified 5 days post-injection (dpi). β1 integrin levels are normalized within each cell line relative to treatment with control siRNA. Fold changes and p values are indicated. Representative images of MDA-MB-231 (C) brain-tropic subclone (231BR) and (D) bone marrow-tropic subclone (231BO) colonization in the zebrafish brain and CVP at 5 dpi. For both cell types, β1 integrin was knocked down prior to cell injection. Arrows indicate cell positions in the brain. Images are average intensity projections from confocal z stacks. The vasculature is displayed in red, the venous circulation in green, and injected cancer cells in blue. Scales are indicated. In the CVP, dashed line shows position of orthogonal view shown in inset. (E) Fraction of cells extravasated in the CVP upon knockdown of β1 integrin. Values were calculated on a per larva basis and normalized to average fraction of control cells extravasated in the CVP for each cell line. **, p<0.01 by Sidak’s multiple comparisons test following two-way ANOVA. N=5 larvae for 231BR Ctrl cells, N=5 larvae for 231BR siβ1 cells, N=5 larvae for 231BO Ctrl cells, and N=5 larvae for 231BO siβ1 cells. Schematic illustrates lack of extravasation in the CVP, with the majority of cells observed at 5 dpi remaining intravascular following knockdown. (F) Fraction of cells extravasated in the brain upon knockdown of β1 integrin. Values were normalized to average fraction of control cells extravasated in the brain for each cell line. ***, p<0.001; ****, p<0.0001 by Sidak’s multiple comparisons test following two-way ANOVA. N=5 larvae for 231BR Ctrl cells, N=5 larvae for 231BR siβ1 cells, N=5 larvae for 231BO Ctrl cells, and N=5 larvae for 231BO siβ1 cells. Schematic illustrates that ~50% of 231BR cells (red) are able to extravasate in the brain following knockdown of β1 integrin, while no extravasation of 231BO cells was observed in the brain upon knockdown. See also Figure S5.

Thus, we reasoned that modulating β1 integrin activity would affect MDA-MB-231 bone marrow-tropic cell homing and initial adhesion in the brain to a greater extent than in brain-tropic cells. Cells were treated with a function-blocking antibody against β1 integrin and injected to the circulation of 2 dpf zebrafish. Intravital imaging was performed from ~4-6 h after injection and continued for the subsequent 12 h (Figure S5C). The cell residence time in each organ and the fraction of cell observed during imaging that remained occluded in the tissue at the end of imaging were calculated. Cells remained largely rounded and within the vasculature immediately following injection for cells treated with both an IgG control and β1 integrin function blocking antibody (Figure S5D-G). Interestingly, modulation of β1 integrin reduced the arrest time of bone marrow-tropic cells at the brain, while little differences were seen in brain-tropic cells (Figure S5H). Moreover, treatment of bone marrow-tropic cells with the function blocking antibody tended to skew arrest of these cells away from the head and towards the CVP compared to control cells treated with an antibody against mouse IgG (Figure S5I). Little difference in cell arrest time or occlusion in the first ~16 h post-injection was observed in the CVP (Figure S5H, I). These results again suggest that bone-marrow tropic cells rely upon β1 integrin function for colonization of the brain to a greater extent that brain-tropic cells.

## DISCUSSION

Tumor cells encounter a myriad of environmental cues along the metastatic cascade before eventual arrest and outgrowth in a secondary organ (Kim and Tanner, 2015; Kumar and Weaver, 2009). While mechanisms of cell escape from a primary tumor have been well characterized (Patsialou et al., 2009; Wyckoff et al., 2007), visualization of the exit from the circulatory system at the latter stages of the cascade are less understood. Moreover, the mechanisms which drive organ selectivity, particularly during early metastasis, are not well characterized. The zebrafish is an excellent model to study dissemination of human tumor cells and eventual colonization in a developmental stage at which the vasculature has properties comparable to tissue properties of mammalian capillary beds. Here, we describe the zebrafish as a tool to begin to address the role of the physical environment on the earliest stages of organotropic metastasis, as we show cells colonize multiple organs and show positive staining for proliferation markers (Figure 1). Intravital imaging of human organotropic cell lines revealed that these cells also displayed non-random colonization of zebrafish organs over a period of 1-5 days after injection (Figure 2). Specifically, human cells that home to murine brain and bone marrow home to analogous organs in the larval zebrafish. Proteomics revealed differential expression of proteins associated with cell mechanosensing machinery between brain- and bone marrow-tropic clones (Figure 3). These differences, concomitant with the differences in vessel architecture in distinct organs, led to selective extravasation. Bone marrow-tropic clones preferentially extravasated from highly disordered vessels in the CVP, while brain-tropic clones preferentially extravasated from linear vessels in the brain (Figure 4). Inhibiting extravasation via knockdown of β1 integrin redistributed observed organ targeting and caused a loss of extravasation in the CVP for both clones (Figure 5). Within a given organ, tissue architecture, composition, mechanics, and regulatory signaling networks provide cues that span length and timescales to drive the emergence of a metastatic lesion (Kim and Tanner, 2015; Oudin and Weaver, 2016). This work emphasizes the importance of the interplay between tissue architecture and cell behavior at the vascular interface to drive preferential colonization of one organ at early stages of metastasis.

Metastatic lesions are often located in organs that are outside the range of single cell imaging techniques. One advantage of the fish is the ability to image the whole organism whilst maintaining single cell resolution not readily achieved in murine systems. In line with Paget’s hypothesis, our results demonstrate that the “soil” contributes to the efficacy of the establishment of a distant lesion. Three pairs of isogenic organotropic clones (MDA-MB-231, 4T1, and BT474m1 derived) displayed non-random targeting in the larval fish (Figure 2). In a recent study, homing of patient derived and cell lines for multiple myeloma to the CVP was observed within a few minutes after injection (Sacco et al., 2016). Here, we observed differential targeting for non-hematological cancers. Moreover, we observe that organ specificity occurred at different times following dissemination, where it can be observed between 1-5 days post-injection. Within this window, cells are able to survive and in a limited capacity proliferate in the presence of the innate immune system. It is worth noting that these results were obtained before adaptive immunity has emerged (Renshaw and Trede, 2012). In the larval fish, brain architecture and cell type are conserved at the developmental stage that was used. However, at this stage, osteoclasts are not fully mature; instead, the CVP is analogous to the hematopoietic stem cell (HSC) niche, both of which contain sinusoidal vascular vessels, fibroblastic reticular cell networks, and perivascular hematopoietic stem and progenitor cells (Kusumbe et al., 2014; Murayama et al., 2006; Nombela-Arrieta et al., 2013; Ramasamy et al., 2016). Moreover, the cytokines implicated in maintaining the niche, such as CXCR4-CXCL12, are conserved between zebrafish and mammals (Ma et al., 2011; Murayama et al., 2006). In the zebrafish model, ~20% of cells injected home to the brain or CVP and are present at 5 dpi. Interestingly, the metastatic efficiency – an indicator of how many cells survive dissemination to exit and colonize – is comparable or greater than what has been observed in murine studies (Fidler, 2003). These results suggest that there is sufficient homology to study initial trafficking steps in multiple organs not readily accessible in some studies. Additionally, in the zebrafish model, ~100 cells are introduced into each fish; thus, it may be possible to screen possible metastatic sites at the time of diagnosis with patient derived tissue.

During metastasis, circulating tumor cells are continuously shed into the circulation system, which is comprised of differential flows and vessel architectures (Kumar and Weaver, 2009). The use of transgenic fish also permits the visualization of key aspects of tumor-capillary dynamics that are not readily created using in vitro mimetics (Figure 3). In the zebrafish system, we are able to begin decoupling the effects of the microenvironment on cell colonization in multiple tissues in vivo within the same animal. Within the fish, a broad diversity of vessel widths and architectures with differential blood flow can be used to interrogate the role of the physical microenvironment. Continuous tracking of multiple cells revealed that cells are move at speeds well below blood flow (Watkins et al., 2012) and comparable to those of zebrafish macrophages (Nguyen-Chi et al., 2015). Interestingly, organotropic MDA-MB-231 cells exhibited similar motility in vivo as has been observed with parental MDA-MB-231 cells within confined environments in an in vitro system (Balzer et al., 2012; Mathieu et al., 2016). Recent work has highlighted the role of physical forces on cell extravasation in the zebrafish CVP (Follain et al., 2018), and we are able to build on these findings by simultaneously examining multiple organs and characterizing vessel architecture and tissue mechanics in detail. In the CVP, vessel architecture is comparable to the tortuosity found within bone marrow, which is distinct from the organized linear blood vessels observed at other organs (Kusumbe et al., 2014; Nombela-Arrieta et al., 2013; Ramasamy et al., 2016). We confirm that the mechanical properties of the organ analogs are also similar to each other on the day of injection, that the tissue mechanics are similar to 3D hydrogel mimetics (Blehm et al., 2016; Kniazeva et al., 2012; Kotlarchyk et al., 2011; Staunton et al., 2016a), and that injected cells can survive in the tissue parenchyma.

Extravasation of cancer cells at secondary sites occurs predominantly within capillary beds and is thought to be a key step in the metastatic cascade to initiate metastatic lesion formation (Nguyen et al., 2009). In this model, extravasation was seen to be the rate limiting step in conveying organ selectivity between brain- and bone marrow-tropic MDA-MB-231 cells. Differences in proteins implicated in mechanosensing were observed for the tumor cell “seeds” (Schwartz, 2010). Of these proteins, β1integrin has been implicated in modulating extravasation for multiple types of tumors using in vivo and in vitro assays (Chen et al., 2016; Stoletov et al., 2010). Here, we show that not only is it differentially regulated between clones, but also that knockdown of β1 integrin inhibits extravasation in an organ- and cell line-dependent manner (Figure 5). In the CVP, extravasation was significantly reduced in both brain- and bone marrow-tropic clones upon knockdown. This finding, in concert with the observation that bone marrow-tropic clones (which have greater endogenous expression of β1 integrin) more efficiently extravasate in the CVP, suggests that β1 integrin-mediated extravasation is a primary mode of cell exit from the bloodstream in that organ. This is in line with previous studies showing that Twist expression regulates β1 integrin-dependent extravasation in the intersegmental vessels (Stoletov et al., 2010). In the brain, knockdown of β1 integrin again inhibits extravasation of bone marrow-tropic cells, but a fraction (~50%) of brain-tropic cells present are still able to extravasate. Function blocking of β1 integrin with an antibody primarily affected cell trafficking and occlusion of the bone marrow-tropic cells in the brain but did not affect brain-tropic cells (**Figure S10**), again suggesting alternative mechanisms for brain-tropic cell colonization of the brain. Previous work has demonstrated that tumor cells in confinement are able to migrate efficiently upon inhibition of β1 integrin activity or cell contractility (Balzer et al., 2012; Stroka et al., 2014), and brain-tropic cells may be utilizing these alternate mechanisms for colonization. Similarly, exosomes implicated in priming distant lesions showed distinct integrin expression patterns for specific organs such as the lung and liver (Hoshino et al., 2015). Interestingly, in that system, exosomes harvested from bone-tropic cells induced vascular leakiness but with a limited repertoire of integrins (Hoshino et al., 2015). Here, our results suggest that organotropic clones may similarly use specific strategies for organ targeting. Taken together, these data suggest that the interplay between tissue architecture and integrin profile may drive non-random extravasation in the zebrafish, and suggest that the larval fish possesses physiological similarities that may be used to screen potential metastatic sites using patient derived cells.

## ACKNOWLEDGEMENTS

This research was supported by the Intramural Research Program of the National Institutes of Health, the National Cancer Institute. The MDA-MB-231 clones were a kind gift of Professor Toshiyuki Yoneda (Indiana University-Purdue University). The Bt474m1 clones were a kind gift from Professor Dihua Yu (Univ. of TX, MD Anderson Cancer Center). The 4T1 clones were a kind gift of Professor Normand Pouliot (Olivia Newton-John Cancer Research Institute, Melbourne, Australia). Transgenic zebrafish lines used for cancer cell injections were kindly provided by Dr. Brant Weinstein (NICHD, NIH). Transgenic zebrafish lines with fluorescent stem cells (CD41) were kindly provided by Blake Carrington and Erica Bresciani (NHGRI, NIH). We thank Dan Castranova, Jian Liu, and Blake Carrington for useful discussions. We would like to thank Susan Garfield and Langston Lim, CCR Confocal Microscopy Core Facility, Laboratory of Cancer Biology and Genetics, NCI for use of the core microscopes.

## AUTHOR CONTRIBUTIONS

K.T., C.P., R.S., N.M. designed experiments, C.P., K.T., K.B. A.D., J.S., E.P. performed experiments, C.P., K.T., L.J., A.D., J.S., S.T. performed data analysis. K.T., C.P., N.M. wrote the main manuscript text. C.P, K.T., J.S. prepared figures. All authors reviewed the manuscript.

## MATERIALS & CORRESPONDENCE

Correspondence and material requests should be addressed to Kandice Tanner, Ph.D., 37 Convent Dr., Bethesda, MD 20852.

## MATERIALS AND METHODS

### Zebrafish husbandry

Animal studies were conducted under protocols approved by the National Cancer Institute and the National Institutes of Health Animal Care and Use Committee. Zebrafish were maintained at 28.5°C on a 14-hour light/10-hour dark cycle according to standard procedures. Larvae were obtained from natural spawning, raised at 28.5°C, and maintained in fish water (60 mg Instant Ocean^©^ sea salt [Instant Ocean, Blacksburg, VA] per liter of DI water). Larvae were checked regularly for normal development. For all experiments, larvae were transferred to fish water supplemented with N-phenylthiourea (PTU; Cat. No. P7629-25G, Millipore Sigma, St. Louis, MO) between 18-22 hours post-fertilization to inhibit melanin formation and maintain optical transparency. PTU water was prepared by dissolving 16 µl of PTU stock (7.5% w/v in DMSO) per 40 ml of fish water. Water was replaced daily. For cancer cell injections, larvae were maintained at 28.5°C until the time of injection (2 dpf) and kept at 33°C following injection. Injected fish were maintained at 33°C until reaching 4 dpf and were then introduced to a larger zebrafish housing system and held in system water at 28.5°C. Regular feeding commenced at 5 dpf. Larvae were collected from the housing system for imaging at 7 dpf.

### Zebrafish lines

For analysis of non-random tissue targeting of MDA-MB-231, Bt474m1, and 4T1 cell lines, the recently described Tg(MRC1a:EGFP/flk:mCherry) zebrafish was used (Jung et al., 2017). In this transgenic strain, EGFP is expressed under the MRC1a promoter to label, at the larval stage, the venous circulation, while mCherry is expressed under the flk promoter to label the vasculature. For analysis of the zebrafish vasculature (vessel size and organization), 2 dpf Tg(fli:EGFP) transgenic fish were used. The Tg(fli:EGFP) strain (Lawson and Weinstein, 2002) was also used in experiments studying the effects of treatment with β1-integrin function blocking antibodies on initial cell trafficking through the vasculature. In experiments for which cells were stained for expression of Ki67, Tg(mpx:EGFP/flk:mCherry) zebrafish, in which neutrophils express EGFP, were used. To visualize hematopoietic stem cell expression in the developing zebrafish larva, transgenic Tg(CD41:EGFP/flk:mCherry) zebrafish, in which CD41-positive cells express EGFP, were used.

### Cell culture

MDA-MB-231 bone marrow-tropic (231BO) and brain-tropic (231BR) cells (Yoneda et al., 2001) were grown and maintained in high glucose (4.5 g/l) DMEM containing 10% FBS, 1% L-glut, and 1% P/S at 37°C. In some cases, cells transformed to express GFP and luciferase (231BR-GFP/Luc) or mKate (231BO-mKate) were used. Bt474m1 and Bt474m1-Br3 cells (Zhang et al., 2013) were maintained in RPMI medium supplemented with 10% FBS, 1% P/S, and 1% L-glut. 4T1Br4 and 4T1BM2 cells (Kim et al., 2018; Kusuma et al., 2012) were maintained in α-MEM containing sodium pyruvate (Cat. No. 10-022-CV, Corning Mediatech, Corning, NY) and supplemented with 5% FBS, 1% P/S, and 1% L-glut. Cells were passaged upon reaching 70-80% confluency, and media was changed every 2-3 days and 24 h before all zebrafish injections. Prior to injection, cells were washed with PBS and detached from the cell culture flask using 10 mM EDTA for 20 min at 37°C. Cells were resuspended in full growth medium, pelleted, resuspended in growth medium again, and counted. Cells were counted, centrifuged at 1000 rpm for 5 min, and resuspended in PBS containing CellTracker Deep Red (Cat. No. C34565, Thermo Fisher Scientific, Waltham, MA) at 1 µM at a concentration of 2×10^6^ cells/ml and incubated at 37°C for 20-30 min. Stained cells were washed twice with PBS, with spins at 2000 rpm for 3 min between washes. After the final spin, cells were resuspended to a concentration of 1 million cells/20 µl in PBS. For integrin-blocking experiments, the final resuspension solution contained 30 µg/ml IgG mouse isotype control (Cat. No. ab91353, abcam, Cambridge, MA) or anti-β1 integrin (abcam, Cat. No. ab24693) antibody. Cells were kept on ice for 1-2 hours during injection, with at least 30 min incubation prior to the first injection.

### siRNA knockdown

Cells were plated at 200,000 cells/well of a 6-well plate in 2.5 ml total growth medium. The following day, cells were transfected with Thermo Fisher Silencer Select siRNA. siRNA was designed to block expression of β1 integrin (Thermo Fisher Scientific, Cat. No. 4390824, siRNA ID s7575) or to be used as a negative control (Silencer Select Negative Control No. 1, Thermo Fisher Scientific, Cat. No. 4390843). siRNA was aliquoted and stored at a concentration of 50 µM and stored at −20°C. Prior to transfection, aliquots were diluted 1:10 in Opti-MEM cell culture medium (Thermo Fisher Scientific, Cat. No. 31985070) to reach a concentration of 5 µM. Per well of a 6-well plate, 3 μl of siRNA at 5 μM was mixed with 150 μl Opti-MEM. In a separate tube, 7.2 μl RNAiMAX (Thermo Fisher Scientific, Cat. No. 13778030) was mixed with 150 μl Opti-MEM. Next, 150 μl diluted of the Opti-MEM/siRNA mixture was combined with 150 μl of the Opti-MEM/RNAiMAX mixture and incubated for 5 min at room temperature. To transfect, 250 µl of this mixture was added to cells plated the previous day, directly to the existing medium. The following day, media was replaced with fresh growth media. Two days following transfection, cell lysate was collected (see below) or cells were used for injection to zebrafish larvae as described above.

### Cell preparation for mass spectrometry

For proteomics analysis of brain- and bone marrow-tropic MDA-MB-231 cells (Figure 3), cells were washed with PBS and detached from cell culture flasks using 10 mM EDTA for 20 min at 37°C. Cells were resuspended in full growth medium, pelleted, resuspended in growth medium again, and counted. Cells were then centrifuged at 1000 rpm for 5 min and resuspended in PBS containing CellTracker Deep Red at 1 µM at a concentration of 2×10^6^ cells/ml and incubated at 37°C for 20 min to match the preparation procedure used for *in vivo* injections. Stained cells were washed twice with PBS, with spins at 2000 rpm for 3 min between washes. After the final spin, cells were resuspended to a concentration of 1 million cells/20 µl in PBS. For the final resuspension, cells were in PBS containing 30 µg/ml IgG. Cells were incubated with the antibody for 30 min), diluted to a concentration of 1e6 cells/ml in serum free media, and seeded in 6-well plate wells (300,000 cells/well). Additional serum free media was added to bring the total volume in each well to 2 ml. Cells were incubated for 8 h at 37°C, 5% CO_2_ prior to lysate collection. This cell preparation method was analogous to cell preparation for injection to the zebrafish. For knockdown experiments, cells were prepared and transfected as described above, and lysate was collected 2 days post-transfection, without staining or detaching cells prior to lysis.

### Mass spectrometry

Lysate was prepared by adding 40 µl/well of lysis buffer containing 1 part Calbiochem 539131 Protease inhibitor cocktail set 1 (Cat. No. 539131, Calbiochem, Millipore Sigma), 1 part phosphatase inhibitor cocktail 2 (Cat. No. P5726, Millipore Sigma), 1 part phosphatase inhibitor cocktail 3 (Cat. No. P0044, Millipore Sigma), and 100 parts M-PER Mammalian Protein Extraction Reagent (Cat. No. 78501, Thermo Fisher Scientific) added. Cells were scraped, and lysate was collected. Lysate was sonicated 20 sec, spun down 5 min at 1600 rpm, and supernatant stored at −80°C prior to mass spectrometry. Lysates were prepared over three separate days to obtain biological triplicates.

Cell lysates (250 μg each) were digested with trypsin using the filter-aided sample preparation (FASP) protocol as previously described with minor modifications (Wisniewski et al., 2009). Lysates were first reduced by incubation with 10 mM DTT at 55 °C for 30 min. Each lysate was then diluted with 8 M urea in 100 mM Tris-HCl (pH 8.5 [UA]) in a Microcon YM-10 filter unit and centrifuged at 14,000 × g for 30 min at 20°C. The lysis buffer was exchanged again by washing with 200 μL UA. The proteins were then alkylated with 50 mM iodoacetamide in UA, first incubated for 6 min at 25 °C and then excess reagent was removed by centrifugation at 14,000 × g for 30 min at 20°C. Proteins were then washed 3 × 100 μL 8 M urea in 100 mM Tris-HCl (pH 8.0) (UB). The remaining urea was diluted to 1 M with 100 mM Tris-HCl pH 8 and then the proteins were digested overnight at 37°C with trypsin at an enzyme to protein ratio of 1:100 w/w. Tryptic peptides were recovered from the filter by first centrifugation at 14,000 × g for 30 min at 20°C followed by washing of the filter with 50 μL 0.5 M NaCl. The peptides were acidified and desalted on a C18 SepPak cartridge (Waters, Milford, MA) and dried by vacuum concentration (Labconco, Kansas City, MO). Dried peptides were fractionated by high pH reversed-phase spin columns (Thermo Fisher Scientific). The peptides from each fraction were lyophilized, and dried peptides were solubilized in 4% acetonitrile and 0.5% formic acid in water for mass spectrometry analysis. Each fraction of each sample was separated on a 75 µm × 15 cm, 2 µm Acclaim PepMap reverse phase column (Thermo Fisher Scientific) using an UltiMate 3000 RSLCnano HPLC (Thermo Fisher Scientific) at a flow rate of 300 nL/min followed by online analysis by tandem mass spectrometry using a Thermo Orbitrap Fusion mass spectrometer. Peptides were eluted into the mass spectrometer using a linear gradient from 96% mobile phase A (0.1% formic acid in water) to 35% mobile phase B (0.1% formic acid in acetonitrile) over 240 minutes. Parent full-scan mass spectra were collected in the Orbitrap mass analyzer set to acquire data at 120,000 FWHM resolution; ions were then isolated in the quadrupole mass filter, fragmented within the HCD cell (HCD normalized energy 32%, stepped ± 3%), and the product ions analyzed in the ion trap.

The mass spectrometry data were analyzed and label-free quantitation performed using MaxQuant version 1.5.7.4 (Cox et al., 2014; Cox and Mann, 2008) with the following parameters: variable modifications - methionine oxidation and N-acetylation of protein N-terminus; static modification – cysteine carbamidomethylation; first search was performed using 20 ppm error and the main search 10 ppm; maximum of two missed cleavages; protein and peptide FDR threshold of 0.01; min unique peptides 1; match between runs; label-free quantitation, with minimal ratio count 2. Proteins were identified using a Uniprot human database from November 2016 (20,072 entries). Statistical analysis was performed using Perseus version 1.5.6.0 (Tyanova et al., 2016). After removal of contaminant and reversed sequences, as well as proteins that were only quantified in one of the three replicate experiments, the label-free quantitation values were base 2 logarithmized and missing values were imputed from a normal distribution of the data. Statistically significant differences were assigned using a two-way t-test with a p-value cut-off of 0.05. Hierarchical clustering was also performed in Perseus. Fold changes were calculated relative to protein expression in 231BR cells from the average base 2 logarithmic differences across biological replicates.

For analysis of signaling pathways that were differentially regulated between the cell types, the annotated list of base 2 logarithmized values and p-values was imported to Ingenuity Pathway Analysis (IPA, QIAGEN, Hilden, Germany). For comparison of 231BR and 231BO cells incubated with the IgG antibody prior to lysate collection, proteins were filtered to retain those with a p-value less than 0.05. This retained 477 of the total 2180 mapped proteins returned from Perseus analysis. Expression pathway analysis was performed in IPA. The top 10 differentially regulated canonical pathways between the cell types (ordered by p value) are listed in Figure 3A. Pathways that involved β1 integrin are displayed in italic text. The activity pattern of each pathway was assessed using the built-in IPA z-score analysis. Pathways are displayed in black if no activity pattern was available.

### Zebrafish circulatory injections

Tricaine stock was prepared by dissolving 400 mg of Tricaine powder (ethyl 3-aminobenzoate methanesulfonate; Cat. No. E10521-50G, Millipore Sigma) with 97.9 ml of deionized water and 2.1 ml of 1 M Tris. Anesthetic fish water was prepared by mixing 4.2 ml of tricaine stock per 100 ml of fish water supplemented with PTU (0.4% buffered tricaine). For injection to the circulation, zebrafish were anesthetized in tricaine and oriented in a lateral orientation on an agarose bed. A volume of 2-5 nL of the cell suspension at 1e6 cells/20 µl (~100-250 cells) were injected directly in the circulation via the posterior cardinal vein, or directly into the hindbrain, using a pulled micropipette. For select experiments, 231BR-GFP/Luc and 231BO-mKate cells were incubated with 30 μg/ml IgG antibody, mixed in equal parts to obtain a total concentration of 1 million cells/20 µl in PBS, and co-injected directly to the zebrafish hindbrain. Injected fish were screened between 4-20 h post-injection to check for successful introduction of cells to the circulatory system or hindbrain and were then imaged or housed for later experiments. Fish lacking cells in the circulation were euthanized.

### Intravital microscopy

Zebrafish larvae were anesthetized using a final concentration of 0.4% buffered tricaine dissolved in fish water that was supplemented with PTU as described above. Anesthetized fish were immobilized in a lateral orientation in 1% (w/v) low gelling temperature agarose (Cat. No. A9414-25G, Millipore Sigma) dissolved in fish water. To enable high-resolution confocal imaging of mounted larvae, fish were laterally oriented in coverglass-bottom chamber slides (Nunc Lab-Tek Chambered #1.0 Borosilicate Coverglass slides, Cat. No 155383, Thermo Fisher Scientific). PTU water supplemented with 0.4% buffered tricaine was then added to the imaging chamber to keep the larvae anesthetized over the course of the experiment.

To acquire images at single-cell resolution within living zebrafish, chamber slides containing the larvae were imaged on Zeiss 710 or 780 laser scanning confocal microscopes. Three-dimensional tile scans of the brain and caudal vein plexus (CVP) were obtained at 1 or 5 dpi to assess cell homing. No overlap was used in tile scans. One-photon, confocal 12-bit images were acquired with a Zeiss 20x EC Plan-Neofluor, 0.3 NA objective and a digital zoom of 1, resulting in a field of view of 425.1 µm × 425.1 µm for each tile of the image. Images were taken at axial steps of 1 µm and stacked to create three dimensional images. Pinhole diameter was set at 90 µm. Samples were simultaneously excited with 488 nm light from an argon laser with a total power of 25 mW, 561 nm light from a solid-state laser with a total power of 20 mW, and 633 nm light from a HeNe633 solid state laser with a total power of 5 mW. Transmitted light was also recorded. Images were taken on two tracks to minimize signal overlap. All lasers were set at or below 8% total power. A beam splitter, MBS 488/561/633, was employed in the emission pathway to delineate the red (band-pass filters ~580-645 nm), green (band-pass filters ~493-574 nm), and far red (band-pass filters ~650-747 nm) channels. The master gain was set at or below 700 for each channel. The zebrafish larva was maintained at 33°C for the course of imaging. Pixel dwell times of 1.58 ms were used.

To image motile cells over the first 12-16 h following injection to the zebrafish circulation, one-photon, confocal, 2-dimensional images were acquired at lateral dimensions of 512 × 512 pixels, which were stacked to acquire 3-dimensional images. Imaging conditions used were:

1. Tg(flk:mCherry/MRC1a:EGFP) or Tg(flk:mCherry/mpx:EGFP) zebrafish were injected with 231BR or 231BO cells stained with Cell Tracker Deep Red prior to injection. Confocal z-stacks were acquired every 10 minutes for ~90 frames at 2 µm axial steps to image a total depth of ~150 µm. 12-bit images were acquired with a Zeiss 20x EC Plan-Neofluor, 0.3 NA objective and a digital zoom of 0.8, resulting in a field of view of 531.37 µm × 531.37 µm. Pinhole diameter was set at 90 µm. Samples were simultaneously excited with 488 nm light from an argon laser with a total power of 25 mW, 561 nm light from a solid-state laser with a total power of 20 mW, and 633 nm light from a HeNe633 solid state laser with a total power of 5 mW. Transmitted light was also recorded. Images were taken on two tracks to minimize signal overlap. All lasers were set at 2% total power. A beam splitter, MBS 488/561/633, was employed in the emission pathway to delineate the red (band-pass filters ~580-645 nm), green (band-pass filters ~493-574 nm), and far red (band-pass filters ~650-747 nm) channels. The master gain was set at or below 700 for each channel. The zebrafish larva was maintained at 33°C for the course of imaging. Pixel dwell times of 1.58 ms were used.
2. Tg(fli:EGFP) zebrafish were injected with 231BR or 231BO cells treated with 30 µg/ml of a function-blocking β1 integrin antibody or IgG control antibody. Cells were stained with CellTracker Deep Red prior to injection. Confocal z-stacks were acquired every 10 minutes for ~90 frames at 2 µm axial steps to image a total depth of ~150 µm. 12-bit images were acquired with a Zeiss Plan-Neofluor 10x/0.30 NA objective and a digital zoom of 1, resulting in a field of view of 850.19 µm × 850.19 µm. Pinhole diameter was set at 100 µm. Samples were simultaneously excited with 488 nm light from an argon laser with a total power of 25 mW and 633 nm light from a HeNe633 solid state laser with a total power of 5 mW. Transmitted light was also recorded. All lasers were set at 2% total power. A beam splitter, MBS 488/633, was employed in the emission pathway to delineate the green (band-pass filters ~493-598 nm) and far red (band-pass filters ~640-747 nm) channels. The master gain was set at or below 700 for each channel. The zebrafish larva was maintained at 33°C for the course of imaging. Pixel dwell times of 1.58 ms were used.

### Zebrafish whole mount immunofluorescence staining

Tg(mpx:EGFP/flk:mCherry) zebrafish were injected at 2 dpi with CellTracker Deep Red stained 231BR or 231BO cells as described above. At 5 dpi, larvae were fixed overnight at 4°C in 4% paraformaldehyde in PBST (PBS supplemented with 0.1% Tween 20). Fixed larvae were washed three times in PBST and permeabilized with 10 µg/ml Proteinase K (Roche, Basel, Switzerland, Cat. No. 03115879001) in PBST for 3 h at room temperature. Larvae were washed four times with PBST and blocked for 2 h at room temperature in PBST containing 5% goat serum. Larvae were then incubated in a 1:50 dilution of anti-Ki67 antibody (Millipore Sigma, Cat. No. AB9260) diluted in PBST with 5% goat serum at 4°C overnight. Stained larvae were washed quickly in PBST four times, and then washed an additional three times in PBST for 10 min each. Fish were blocked in 5% goat serum in PBST for 2 h at room temperature prior to secondary antibody addition. Larvae were transferred to a 1:200 (10 µg/ml) secondary antibody cocktail containing an AlexaFluor 405 goat anti-rabbit secondary antibody (Thermo Fisher Scientific, Cat. No. A31556) in PBST with 5% goat serum. Fish were stained for 3 h at room temperature, washed three times in PBST, and imaged. Presented images are average intensity projections of confocal z slices containing cells taken at 20x magnification. Look up tables were adjusted to minimize background.

### Quantification of vessels size and topography of the zebrafish vasculature

Whole-larva tile scans (15% tile overlap) of 2 dpf Tg(fli:EGFP) zebrafish were acquired on a Zeiss 780 LSM confocal microscope with a 20x Zeiss 20x EC Plan-Neofluor, 0.3 NA objective and a digital zoom of 1 using an axial step of 1 µm. Average intensity projections of the entire larvae were made in Fiji. The line tool in Fiji was used to obtain widths of 10 vessels per fish in four fish in the brain, CVP, and ISV for a total of N=40 vessels at each location. To characterize vessel topographical complexity, the circle tool was used to draw 10, 20, 50, or 100 µm-diameter circular regions of interest. This region of interest was randomly placed over vessels in the brain, CVP, or ISV, and the Orientation J plugin in Fiji was used to measure the coherency (defined here as the order parameter) at each region of interest and is described in detail elsewhere (Fonck et al., 2009; Puspoki et al., 2016). The order parameter ranged from a value of 0 for isotropic distributions to 1 for aligned regions. For each region of interest size and tissue (brain, CVP, or ISV), 10 regions were measured for 4 fish, giving a total number of N=40 regions.

### Quantification of cell numbers and extravasated cells in the zebrafish brain and CVP

Fish injected with cells were screened for health and successful dissemination of cells throughout the circulatory system prior to incubation at 33°C (2-4 dpf) and 28.5°C (4-7 dpf). At 5 dpi, fish were randomly selected from the husbandry tank, anesthetized, mounted in agarose, and imaged as described above.

Cells were counted as being in the CVP if they were within or proximal to the cardinal artery and caudal vein plexus, from the first 15 intersegmental vessels to the end of the cardinal artery. Cells were counted as being in the brain if they were within the first 700 µm from the most distal location of arteries in the head. Cells in the heart sac, Duct of Cuvier, and aortic arches were not counted as being in the brain. In the CVP, cells that were present at the injection site were excluded from analysis. The minimum size of particles in the deep red channel to be counted as a cell was 10 μm × 10 μm, as determined by the size of the particle at the z slice of its largest diameter in Fiji. Cells were counted as extravasated if they were outside of the vessel lumen, judged by 3D volume reconstructions in Fiji. Cells were counted from: N=21 fish for 231BR cell injections; N=23 fish for 231BO cell injections; N=28 fish for 4T1Br4 cell injections; N=25 fish for 4T1BM2 cell injections; N=7 fish for Bt474m1-Br3 cell injections; and N=11 fish for Bt474m1 cell injections. Cells were counted in larvae at 5 dpi (231BR, 231BO, 4T1Br4, 4T1BM2) or 1 dpi (Bt474m1-Br3, Bt474m1).

To determine whether cells homed to the brain or CVP, the total number of cells (including intravascular and extravascular cells) in the brain and CVP as described above were counted for each larva. For a given larva, the ratio of cells in the brain to the total number of cells counted in the brain and CVP was calculated. Cells were scored as targeting the brain in a given larva if this ratio was greater than 0.3 and as homing to the CVP when this ratio was less than or equal to 0.3. The distribution of larvae scored as brain or CVP homing for each cell line is summarized in Table 1. All fish at 5 dpi with at least one cell present were included in analysis. Trafficking of brain-targeting vs. bone-targeting or parental cells for the analysis of cell homing (Figure 2E-I) was statistically analyzed using a two-tailed binomial test implemented in GraphPad Prism, with the expected percentage of larvae exhibiting targeting to a particular tissue set by the bone marrow-targeting (231BO, 4T1BM2) or parental (Bt474m1) cell lines.

**Table 1.** Summary of cell homing in zebrafish larvae

| Cell line | $\frac{\text{Cells in brain}}{\text{Cells in (brain + CVP)}} \leq 0.3$ | $\frac{\text{Cells in brain}}{\text{Cells in (brain + CVP)}} > 0.3$ |
| --- | --- | --- |
| 231BR | 10 | 11 |
| 231BO | 20 | 3 |
| 4T1Br4 | 20 | 8 |
| 4T1BM2 | 20 | 5 |
| Bt474m1-Br3 | 1 | 6 |
| Bt474m1 | 7 | 4 |

To examine 4T1 cell extravasation in the brain, the subset of cells present in the brain tissue that had extravasated were counted for each larva. The total number of larvae from the population described above with no cells extravasated in the brain or 2 or more cells extravasation in the brain were counted. This sample encompassed larvae from N=21 fish for 4T1Br4 cell injections, and N=23 fish for 4T1BM2 cell injections. The remainder of fish had one cell extravasated in the brain and were excluded from further analysis for Figure 2J. In Figure 2J, the fraction of larvae scored in each extravasation group is plotted. Extravasation in the brain was statistically analyzed using a two-tailed binomial test implemented in GraphPad Prism, with the expected percentage of larvae exhibiting no extravasation or more than 2 cells extravasated set by the bone marrow-targeting 4T1BM2 line.

**Table 2.** Summary of 4T1 cell extravasation in the brain

| Cell line | 0 cells extravasated in brain | $\geq 2$ extravasated in brain |
| --- | --- | --- |
| 4T1Br4 | 13 | 8 |
| 4T1BM2 | 19 | 4 |

To study topography-mediated extravasation of MDA-MB-231 cells in the brain and CVP, the number of cells that had extravasated in the brain and CVP were counted for each larva. For a given larva, the ratio of cells extravasated in the brain to the total number of cells extravasated in the brain and CVP was calculated. Zebrafish in which no extravasation had occurred in either organ were excluded from this analysis. This sample encompassed larvae from N=21 fish for 231BR cell injections, and N=22 fish for 231BO cell injections. The population average extravasation ratio between isogenic clones was analyzed using Welch’s t test.

Finally, to study the effect of siRNA-mediated knockdown of β1 integrin on cell survival and extravasation, the total number of control and knockdown cells homing to and extravasated in the brain and CVP were counted as described above. The total number of cells present was the sum of cells homing to the brain and CVP and was counted for: 231BR Ctrl siRNA – 5 embryos, 48 total cells; 231BR β1 siRNA – 5 embryos, 15 total cells; 231BO Ctrl siRNA – 5 embryos, 33 total cells; 231BO β1 siRNA – 5 embryos, 8 total cells. The total cell count was averaged and normalized to the average of Ctrl siRNA-transfected cells for both 231BR and 231BO cells. Averages were compared by two-way ANOVA with Sidak’s multiple comparisons post-test between Ctrl and siβ1 transfected cells for each cell line. Cell homing to the brain and CVP was quantified using the ratio of cells present in the brain to cells present in the brain and CVP, and this fraction was again normalized against the average value for Ctrl siRNA-transfected cells for both 231BR and 231BO cells. Averages were compared by two-way ANOVA with Sidak’s multiple comparisons post-test between Ctrl and siβ1 transfected cells for each cell line. Within both the brain and CVP, the fraction of cells present in the organ that were extravasated was calculated on a per larva basis, and averages were normalized to the average value for the same cell type transfected with Ctrl siRNA for that organ. The number of larvae used in this calculation for each organ is summarized below. Within each organ, averages were compared by two-way ANOVA with Sidak’s multiple comparisons post-test between Ctrl and siβ1 transfected cells for each cell line.

**Table 3.** Summary of animals used in β1 integrin knockdown studies

| Cell type | siRNA | Organ | Number of larvae |
| --- | --- | --- | --- |
| 231BR | Ctrl | CVP | 5 |
| 231BR | $\beta 1$ integrin | CVP | 5 |
| 231BO | Ctrl | CVP | 5 |
| 231BO | $\beta$ 1 integrin | CVP | 5 |
| 231BR | Ctrl | Brain | 5 |
| 231BR | $\beta$ 1 integrin | Brain | 5 |
| 231BO | Ctrl | Brain | 5 |
| 231BO | $\beta$ 1 integrin | Brain | 5 |

### Quantification of zebrafish neutrophil accumulation in the head and tail

Tg(flk:mCherry/mpx:EGFP) zebrafish were injected with MDA-MB-231 brain-tropic or bone marrow-tropic cells at 2 dpf. Larvae were screened for successful cell dissemination through the circulatory system, mounted in agarose, and imaged at 1 dpi as described above. Images were oriented along the horizontal axis and then cropped. Images in the zebrafish tail were cropped vertically to a height of 512 pixels (425.10 µm) and horizontally to encompass the tail and 15 most posterior ISVs in Fiji. Images of the zebrafish head were cropped to a size of 844×722 pixels (width × height; 700.74 µm × 599.45 µm) from the most anterior blood vessel in the head. Confocal z stacks were maximum intensity projected, and the channel showing zebrafish neutrophils was processed in Fiji. Images were auto threshold using the Huang method and automatic Fiji binarization threshold. Binarized images were run through the Fiji watershed algorithm, and the number of particles present between the sizes of 20-10000 µm^2^ were counted. Larvae in which background fluorescence intensity was too high to obtain accurate counts were discarded from further analysis. Average neutrophil counts in the head and tail were calculated for N=7 larvae injected with brain-tropic cells (total of 255 neutrophils counted in the head and 421 neutrophils counted in the tail) and N=5 larvae injected with bone-tropic cells (total of 360 neutrophils counted in the head and 490 neutrophils counted in the tail). Neutrophil numbers were analyzed by two-way ANOVA with Tukey’s multiple comparisons post-test. For this population, the average ratio of neutrophils in the head to total neutrophils counted in the head and tail was also calculated on a per larva basis. This ratio was analyzed between larvae injected with brain- and bone marrow-tropic cells using a two-tailed t test.

### Intravital cell tracking

Time-lapse microscopy images were exported to Fiji for analysis. To adjust for fish growth during imaging, images were first registered using the Correct 3D Drift plugin (Parslow et al., 2014), with the vasculature of the fish used as a topographical reference. For registration, the multi-time scale computation, subpixel drift correction, and edge enhancement options were enabled. Cancer cells were tracked in 3D in registered images over a 12 h imaging window from the registered image stacks using the TrackMate plugin for Fiji (Tinevez et al., 2017). Images were segmented in the cancer cell fluorescence channel in each frame using a Laplacian of Gaussian (LoG) detector with a 15 µm estimated particle diameter. An initial threshold of 1.0 was set, and the sub-pixel localization and median filter options in TrackMate were activated. Segmentation was then further refined by manual adjustment of the threshold to minimize detection of background objects. Segmented objects were linked from frame to frame with a Linear Assignment Problem (LAP) tracker with 30 µm maximum frame-to-frame linking distance. Tracks were visually inspected for completeness and accuracy over the entire acquisition period and were manually edited to ensure that point-to-point tracks were generated for the entire time that a cell was in the field of view.

### Calculation of cell speed and residence time in the zebrafish

Cell tracks (x,y,t,z) were exported to MATLAB for analysis. For each cell, frame-to-frame speed was calculated by dividing the displacement of the cell by the time interval between frames. An average speed for that cell over the course of imaging was then calculated by averaging these frame-to-frame speeds, and that average speed was used in subsequent plots and calculations. Speeds were only calculated for those cells remaining in frame for at least 4 frames (30 min). Speeds plotted in Figure 3 are grouped measurements (231BR – 91 cells in brain, 131 cells in CVP; 231BO – 46 cells in brain, 56 cells in CVP) made from multiple fish (231BR – 6 larvae; 231BO – 8 larvae). Speeds were compared by two-way ANOVA with Sidak’s multiple comparisons post-test between cell types in each tissue. This is the same method used for tracking cells in the intersegmental vessels in vivo, where cells were tracked in the ~12 h following injection to the circulation of 2 dpf Tg(fli:EGFP) zebrafish (231BO – 14 cells in 7 larvae; 231BR – 38 cells in 6 larvae).

To calculate residence time in the brain and CVP upon function blocking of β1 integrin, the number of frames that cells were tracked were simply counted and multiplied by the imaging interval (10 min) for observations made during the total analysis period, which was set at 72 frames. Cells that were in frame for only one frame were not considered. Cells were grouped across conditions to obtain population average residence times for a given condition. The average residence time for cells treated with the function blocking antibody was divided by corresponding values for cells treated with the IgG isotype control antibody. Normalized residence times upon function blocking were statistically analyzed between organs for a given cell type using Sidak’s multiple comparisons post-test following two-way ANOVA. To estimate the percentage of cells becoming occluded in each tissue over the first 72 frames imaged, the number of cells present at the final frame (12 h) was divided by the total number of cells observed in all fish larvae. In each organ, this fraction was divided by the fraction observed for cells treated with the control IgG antibody. Cells that were in frame for only one frame were not considered. Residence times and occlusion fractions were calculated from: 231BR IgG – 5 larvae, 207 cells in head, 159 cells in CVP; 231BR anti-β1 – 4 larvae, 181 cells in head, 296 cells in CVP; 231BO IgG – 5 larvae, 78 cells in head, 139 cells in CVP; anti-β1 – 3 larvae, 26 cells in head, 77 cells in CVP.

### Microchannel experiments

Master molds for the microchannel devices were fabricated from SU-8 on silicon wafers using standard photolithographic techniques for dual-height structures. Two chrome-glass contact masks were designed using L-Edit (Mentor Graphics, Wilsonville OR) and purchased from Front Range Photomask (Lake Havasu City, AZ). Four-inch silicon wafers (NOVA Electronic Materials, Flower Mound, TX) were first cleaned with acetone, isopropanol, and water, then dehydrated for 10 minutes on a 200°C hotplate. Three layers of SU-8 photoresist (Microchem, Newton MA) were used to pattern the devices, with minor modifications of the manufacturer suggested protocols. First, an adhesion layer of SU-8 2000.5 was spun on the wafer using a single wafer spin coater (Model WS-650S-8NPP/LITE, Laurell Technologies, North Wales, PA), softbaked, flood-exposed with the mask aligner (Model 208, Optical Associates, Inc., San Jose, CA), and post-exposure baked. Second, the microchannel layer was patterned using the first chrome-glass mask with SU-8 2005 to make an array of 7 μm high, 8 μm wide channels; the SU-8 2005 layer was softbaked, exposed, and post-exposure baked. Both the SU-8 2000.5 and the SU-8 2005 were prepared in our laboratory by dilution of SU-8 2050 with SU-8 thinner (Microchem) to obtain the desired percent solids as per the manufacturer product sheets.

The large fluid channels were patterned with SU-8 2100 as received, using the second chrome-glass mask. Solvent edge bead removal was performed during the spinning step to ensure good contact between the mask and resist, and alignment marks were used to ensure the small microchannels were centered in the gap of the larger channels. For the thick layer, a long-pass filter (PL-360-LP, Omega Optical, Brattleboro, VT) was used during exposure to prevent overexposure of the top of the resist layer. After this final layer was softbaked, exposed, and post-exposure baked, the pattern was developed with SU-8 developer, rinsed thoroughly, and hardbaked at 150°C for two minutes. The templates were then treated with silane vapor (tridecafluoro-1,1,2,2-tetrahydrooctyl trichlorosilane (Gelest, Morrisville, PA) for one hour to reduce adhesion during use.

Final microfluidic devices were formed using replica molding with polydimethylsiloxane (PDMS; Sylgard 184 kit, Dow Corning, Midland, MI, USA) at a ratio of 10 parts prepolymer to 1 part crosslinker. The PDMS components were mixed, poured over the patterned wafer, and degassed. PDMS devices were polymerized at 85°C for ~2 h before removal from the master mold. Devices were then cut, and fluid access ports were punched with 4 mm biopsy punches (Miltex 69031-05, Electron Microscopy Sciences, Hatflield, PA).

PDMS devices were cleaned with ethanol and DI water. Devices and two-well coverglass-bottom chamber slides (Nunc Lab-Tek Chambered #1.0 Borosilicate Coverglass slides; Cat. No 155380, ThermoFisher Scientific) were exposed to oxygen plasma for 30 sec. PDMS devices were then irreversibly bound to the slides and filled with PBS. Device walls were coated with 20 μg/ml rat tail collagen type I, 10 μg/ml fibronectin from human plasma (Millipore Sigma, Cat. No. FC010), or 1% BSA (all dissolved in PBS) by adsorption for 1 h at 37°C, 5% CO_2_. Devices were washed with PBS following removal of the protein solution.

231BR and 231BO cells transformed to express green fluorescent protein (GFP) and luciferase (Luc), hereafter referred to as 231BR-GFP/Luc and 231BO-GFP/Luc, respectively, were washed with PBS, detached with 0.25% trypsin-EDTA, and resuspended in serum containing medium at 1×10^6^ cells/ml. Cells were introduced to the device by adding 20 µl of the cell seeding solution to the device inlet well, and the device was placed in the incubator for 15 min for cell attachment. If cell inlet flow stopped during this period, some of the seeding solution was moved to the other cell well to re-initiate flow. Following seeding, devices were washed by adding serum free medium to each well and incubating the device for 30 min. Devices were then washed three times using medium with serum. After the final wash, medium with serum was added to all wells, with an addition 2.5 ml/well of serum containing medium added to each well of the chamber slide to keep the device immersed and minimize evaporation during imaging. Thus, cells migrated in the device in the absence of flow and chemical gradients.

To obtain live cell images of migrating cells, devices were mounted in a stage top incubator on a Zeiss 780 LSM confocal microscope ~2 h after the final device wash. Devices were maintained at 37°C, 5% CO_2_ and imaged every 10 min for up to 12 h using a Zeiss Plan-Neofluor 10x/0.30 NA objective and a digital zoom of 1, resulting in a field of view of 850.19 µm × 850.19 µm. Multiple positions were imaged to obtain fields of view encompassing the entire device. Samples were excited with 488 nm light from an argon laser with a total power of 25 mW to capture 2048 pixel × 2048 pixel, 16 bit, 2 line mean averaged images with the pinhole width set at 90 µm. The laser was set at 2% of total power. Biological duplicates were run for each combination of cell type and ECM protein, with the exception of BSA, where three devices were analyzed for each cell type to obtain a sufficient number of cells in microchannels.

### Calculation of cell speed from microchannel experiments

Migrating cells were tracked within the microchannels. To track cells, the GFP channel of each image was first preprocessed to better visualize cell morphology. In Fiji, the background was subtracted using the rolling ball method with a radius of 50 pixels. Images were then subjected to a Gaussian blur with sigma=2 pixels, following by an unsharp mask at a radius of 2 pixels and a mask value of 0.60. Processed images were saved as tiff files and tracked using TrackMate (Tinevez et al., 2017). Images were segmented using the Downsample LoG detector with a 40 µm estimated particle diameter, threshold of 0.1, and downsampling factor of 2. The entrance and exit of the microchannels was manually identified, and tracking was confined to either the regions where cells were fully within channels. Only tracks containing at least 12 points were retained for analysis. After initial track assignment using a LAP tracker, tracks were manually checked and updated for completeness and accuracy. Tracking was discontinued if a cell reached the end of a microchannel or exited the base of a microchannel, encountered a cell in the channel traveling in the opposite direction, or was in contact with a preceding cell in the microchannel.

Cell tracks (x,y,t,z) were exported to MATLAB for analysis. For each combination of cell type and ECM protein, 30 tracks from each biological replicate were randomly selected, and an average speed for a given cell was obtained by averaging the frame-to-frame speed (calculated by dividing the displacement of the cell by the time interval between frames) over the first 30 frames for which the cell was tracked. For BSA coated devices, fewer cells entered the microchannels, and fewer than 30 cells were used per biological repeat in some instances. Average speeds were grouped from each biological replicate to obtain an ensemble of cell speeds. This ensemble was averaged to obtain a mean cell migration speed for a given condition. Average speeds between cell lines for a given ECM coating and compared to migration in the ISV were analyzed by two-way ANOVA with Sidak’s multiple comparisons post-test in GraphPad Prism 7.

**Table 4.** Summary of cells tracked in confined migration experiments.

| Cell type | Protein coating | Biological replicates | Total number of cells tracked |
| --- | --- | --- | --- |
| 231BR | Collagen type I | 2 | 60 |
| 231BO | Collagen type I | 2 | 60 |
| 231BR | Fibronectin | 2 | 60 |
| 231BO | Fibronectin | 2 | 60 |
| 231BR | BSA | 3 | 52 |
| 231BO | BSA | 3 | 57 |

### Optical tweezer-based microrheology

For all experiments, at 24 hours post fertilization (hpf), larvae were transferred to fish water supplemented with PTU to inhibit melanin formation for increased optical transparency. Larvae were then returned to the incubator at 28.5°C and checked for normal development. Zebrafish larvae at 2 dpf were anesthetized using 0.4% buffered tricaine. Polystyrene beads, 1 micron in diameter resuspended in PBS with a final cell concentration of 1 × 10^7^ beads/ml, were injected directly into the organ of interest as previously described (Blehm et al., 2016; Staunton et al., 2017). Larvae were then anesthetized using 0.4% buffered tricaine, then embedded in a lateral orientation in 1% low melting point agarose and allowed to polymerize in with cover glass (no. 1.5 thickness) as previously described (Blehm et al., 2016). Egg water supplemented with tricaine was added to the agarose hydrogel for the entire time of data acquisition. For complete experimental details, see (Blehm et al., 2016; Staunton et al., 2017; Staunton et al., 2016b).

Our home-built setup consists of a 1064 nm trapping beam steered by an acousto-optic deflector to oscillate the trap and a stationary 975 nm detection beam that is coupled into and colocated with the trap with a dichroic before being sent into the backport of an inverted microscope with a long working distance water objective and a high NA condenser. Telescope lenses conjugate the optical plane at the acousto-optic deflector (AOD) to the back aperture of the condenser, which is placed in Kohler illumination after the object is focused in the specimen place. Above the condenser, the detection beam is relayed to a quadrant photodiode for back focal plane interferometric position detection. Each bead is positioned precisely in the center of the trap by scanning it through the detection beam in three dimensions using a piezo nanopositioning stage while recording the voltages from the QPD. The V-nm relation of the QPD is calibrated in situ by fitting the central linear region of the detector response to scanning the bead through the detection beam in the direction of the oscillations, giving β in V/nm. A second QPD records the position of the trapping laser to find the relative phase lag between the bead and trap oscillations. The optical trap stiffness k is determined in situ from the thermal power spectrum of each measured bead while the trap is stationary, using the active-passive calibration method (Fischer and Berg-Sørensen, 2007). Together with β, k, and the bead’s mass m and radius a, the trajectories yield the complex modulus as a function of frequency, G*(ω), of each bead’s surrounding microenvironment. In this equation, the complex modulus, G*(ω), can be broken down into components, with *G*\*(*ω*)=*G*′(*ω*)+*iG*″(*ω*), where the real part, G’(ω), is the elastic component and the imaginary part, G’’(ω), is the viscous component. The complex modulus, G*(ω), is calculated as 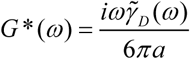, where the friction relaxation spectrum 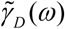 is related by the equation 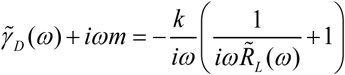 to the active power spectrum 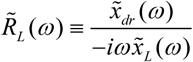, with 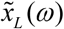 and 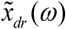 the Fourier transforms of the time series of the positions of the trapping laser and the driven bead respectively, recorded while the trap is oscillating. The stiffness 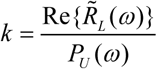 is determined from the real part of the active power spectrum and the passive power spectrum, 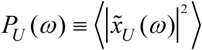, where 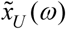 is the Fourier transform of the time series of the undriven bead’s thermally fluctuating position while the trap is held stationary. Each bead is subjected to fourteen consecutive 2s pulses, with the trap alternately oscillating or stationary. Amplitude of oscillations was set to 20nm with power of 100mW at the back aperture. Only probes at distances exceeding ~30 μm away from the cover slip surface to minimize drag in consideration of Faxen’s law were measured (Neuman and Block, 2004).

Samples were measured in triplicate with at least 30 beads per sample measured. Laser power was set to 100mW at the back aperture. Data were analyzed using custom MATLAB programs. Experiments were controlled using custom LabVIEW programs. Active microrheology data are presented as complex modulus (mean ± standard deviation), normalized to the complex modulus of the CVP at 2 dpf. Data are also represented as a function of frequency from 3Hz to 15kHz. Data normalized to the complex modulus of the CVP were analyzed by two-way ANOVA with Tukey’s honestly significant difference post-test.

---

**Figure S1.**
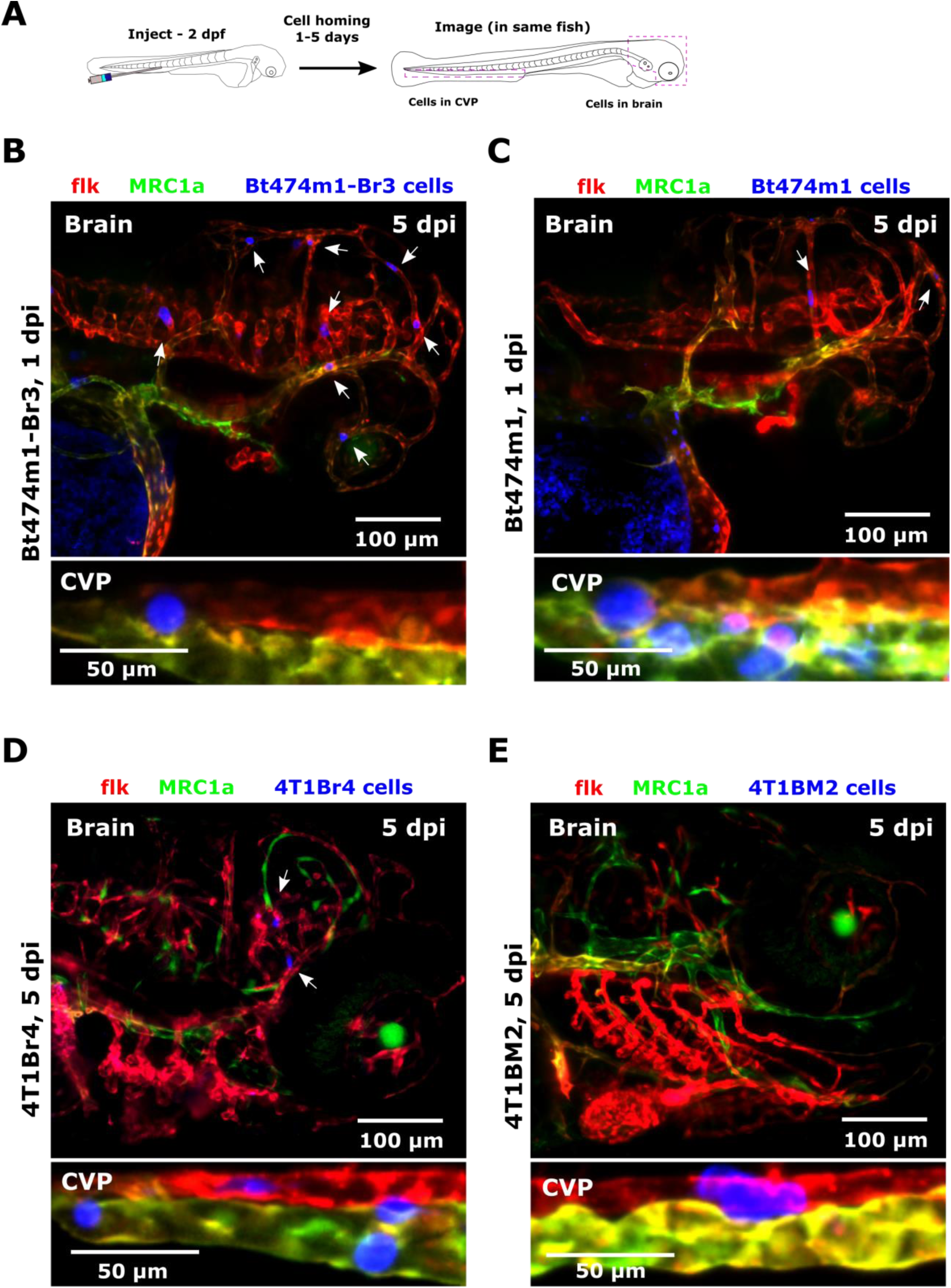
Related to Figure 2. Non-random targeting of organ-tropic human cell lines in a zebrafish metastasis model. (A) Schematic of experimental setup. Cancer cells were injected into the circulatory system of Tg(flk:mCherry/MRC1a:EGFP) zebrafish at 2 days post-fertilization (dpf) and imaged at 1–5 days post-injection (dpi; zebrafish age at imaging was 7 dpf). Confocal z stacks were acquired in the brain and caudal vein plexus (CVP) of each larva. Representative images of Bt474m1 (A) brain-tropic cells (Bt474m1-Br3) and (B) parental cells (Bt474m1) at 1 dpi in the brain and CVP. Representative images of 4T1 (A) brain-tropic cells (4T1Br4) and (B) bone marrow-tropic cells (4T1BM2) at 5 dpi in the brain and CVP. Arrows indicate cell positions in the brain. Scales are indicated. See also Figure 2.

**Figure S2.**
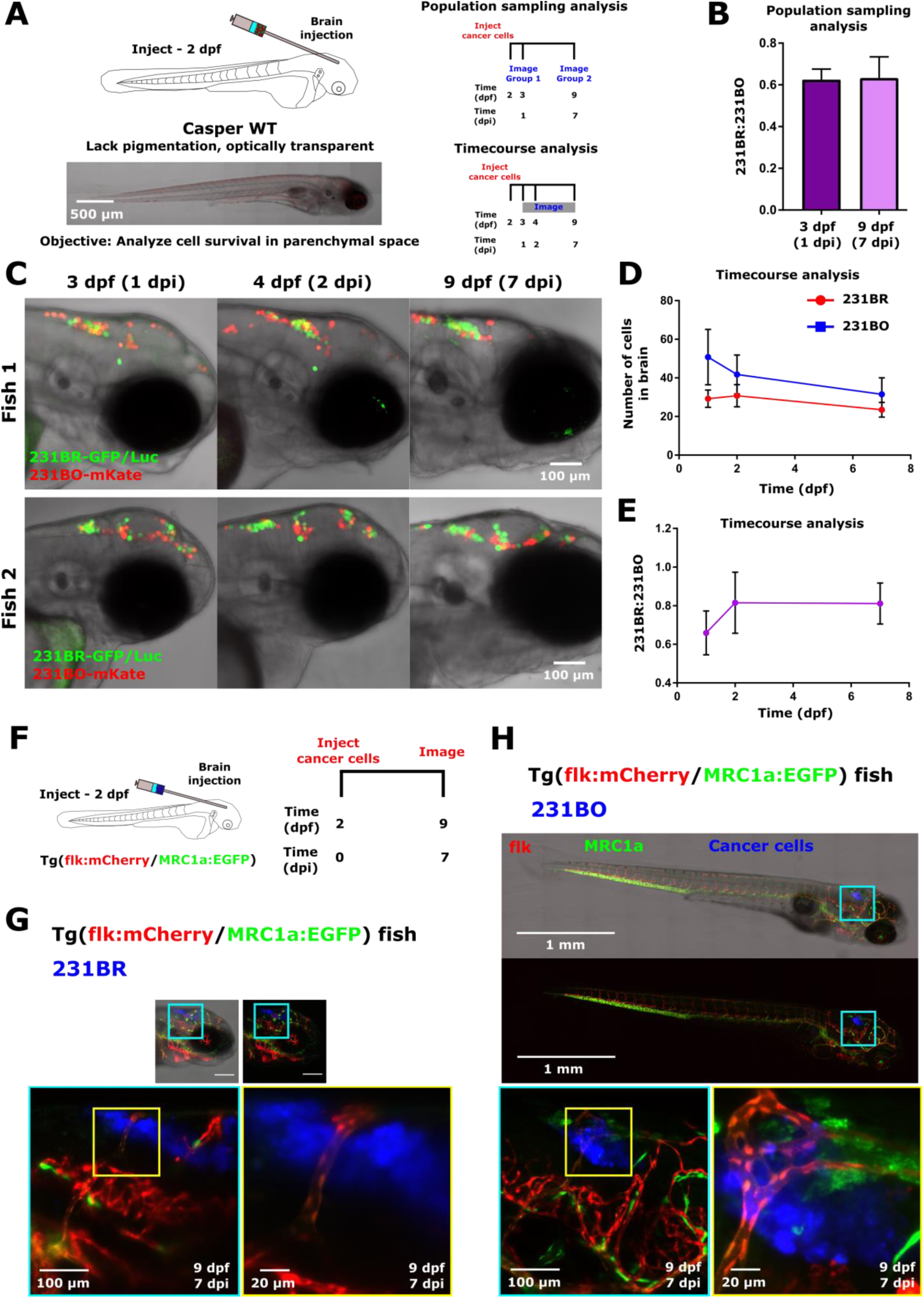
Related to Figure 2. MDA-MB-231 brain-tropic and bone marrow-tropic cell lines exhibit equal levels of survival following direct injection to the brain parenchyma. (A) Schematic of direct brain injection experiments. Brain-tropic MDA-MB-231 cells stably expressing GFP and luciferase (231BR-GFP/Luc) and bone marrow-tropic MDA-MB-231 cells stably expressing mKate (231BO-mKate) were co-injected into the brain of non-pigmented Casper WT zebrafish at 2 days post-fertilization (dpf). A representative image of the fish at 7 days post-injection (dpi; zebrafish is 9 dpf) is shown for reference. Cell survival was assessed by two methods. For population sampling of co-injected cells, a subset of fish was imaged at 1 dpi prior to being euthanized. At 7 dpi, a second subsampling from the injected pool of fish was imaged. Alternatively, for timecourse analysis, injected larvae were imaged at 1, 2, and 7 dpi, and total cell numbers were counted on each day. (B) Quantification of the ratio of 231BR:231BO cells present from population subsamplings at 1 dpi and 7 dpi. Plots show mean ± SEM. N=10 fish at 1 dpi (520 total cells counted), N=6 fish at 7 dpi (142 total cells counted). Difference was not significant by unpaired t test. (C) Two representative examples of cell survival following direct injection into the brain parenchyma. Each zebrafish is imaged at 1, 2, and 7 dpi. 231BR-GFP/Luc cells are shown in green, and 231BO-mKate cells are displayed in red. Scales are indicated. (D) Numbers of 231BR-GFP/Luc and 231BO-mKate cells present over time following co-injection to the brain. Plot displays mean ± SEM cells from N=4 injected zebrafish. (E) Quantification of the ratio of 231BR:231BO cells present over time following co-injection to the brain. Plot displays mean ± SEM cells from N=4 injected zebrafish. (F) Schematic of single cell type injection experiments. MDA-MB-231 brain-tropic (231BR) or bone marrow-tropic (231BO) were stained and, injected directly to the brain of 2 dpf Tg(flk:mCherry/MRC1a:EGFP zebrafish), and imaged at 7 dpi. Representative images of (G) 231BR and (H) 231BO cells in the brain at 7 dpi are shown. Images are average intensity projections from confocal z stacks. Inset locations are indicated by the colored boxes. The vasculature is displayed in red, the venous circulation in green, and injected cancer cells in blue. Scale bars in top panels (overview images) are 1 mm. Scales are indicated in inset images. See also Figure 2.

**Figure S3.**
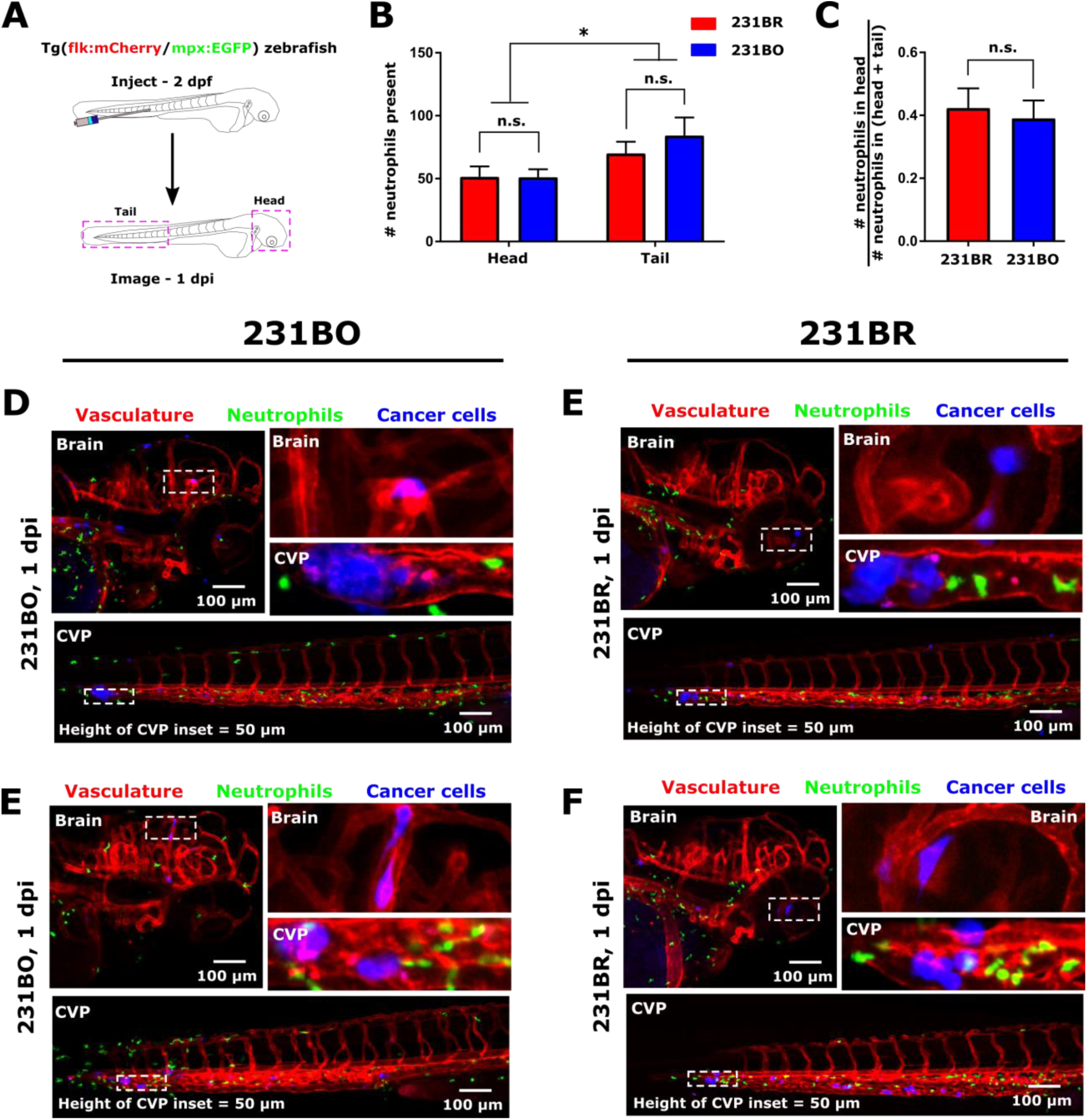
Related to Figure 2. Zebrafish neutrophils respond similarly to organ-tropic cell line injection. (A) Schematic of experiment. Brain-tropic MDA-MB-231 cells (231BR) or bone marrow-tropic MDA-MB-231 cells (231BO) were injected into the circulation of 2 days post-fertilization (dpf) Tg(flk:mCherry/mpx:EGFP) zebrafish. The zebrafish head and tail were imaged at 1 day post-injection (dpi), when zebrafish were 3 dpf. (B) Average ± SEM of zebrafish neutrophil counts in the head and tail 1 day after injection with 231BR or 231BO cells. Tissue of interest but not cell type was a significant source of variation by two-way ANOVA (*, p<0.05). Neutrophil counts for each cell type and region were not significantly different by Tukey’s multiple comparisons test. N=7 larvae injected with 231BR cells, N=5 larvae injected with 231BO cells. (C) Average ± SEM of the ratio of neutrophils present in the head to neutrophils present in the head and CVP. Values were calculated on a per larva basis. n.s., not significant by two-tailed t test. N=7 larvae injected with 231BR cells (total of 255 neutrophils counted in the head and 421 neutrophils counted in the tail), N=5 larvae injected with 231BO cells (total of 360 neutrophils counted in the head and 490 neutrophils counted in the tail). Representative images of zebrafish injected with (D,E) bone marrow-tropic and (F,G) brain-tropic MDA-MB-231 cells. Images display average intensity projections from confocal z stacks. Blood vessels are displayed in red, zebrafish neutrophils in green, and human cancer cells in blue. Inset locations are indicated by the dashed white boxes. For scale, the height of the CVP inset = 50 μm. Two representative images for each cell line are shown. See also Figure 2.

**Figure S4.**
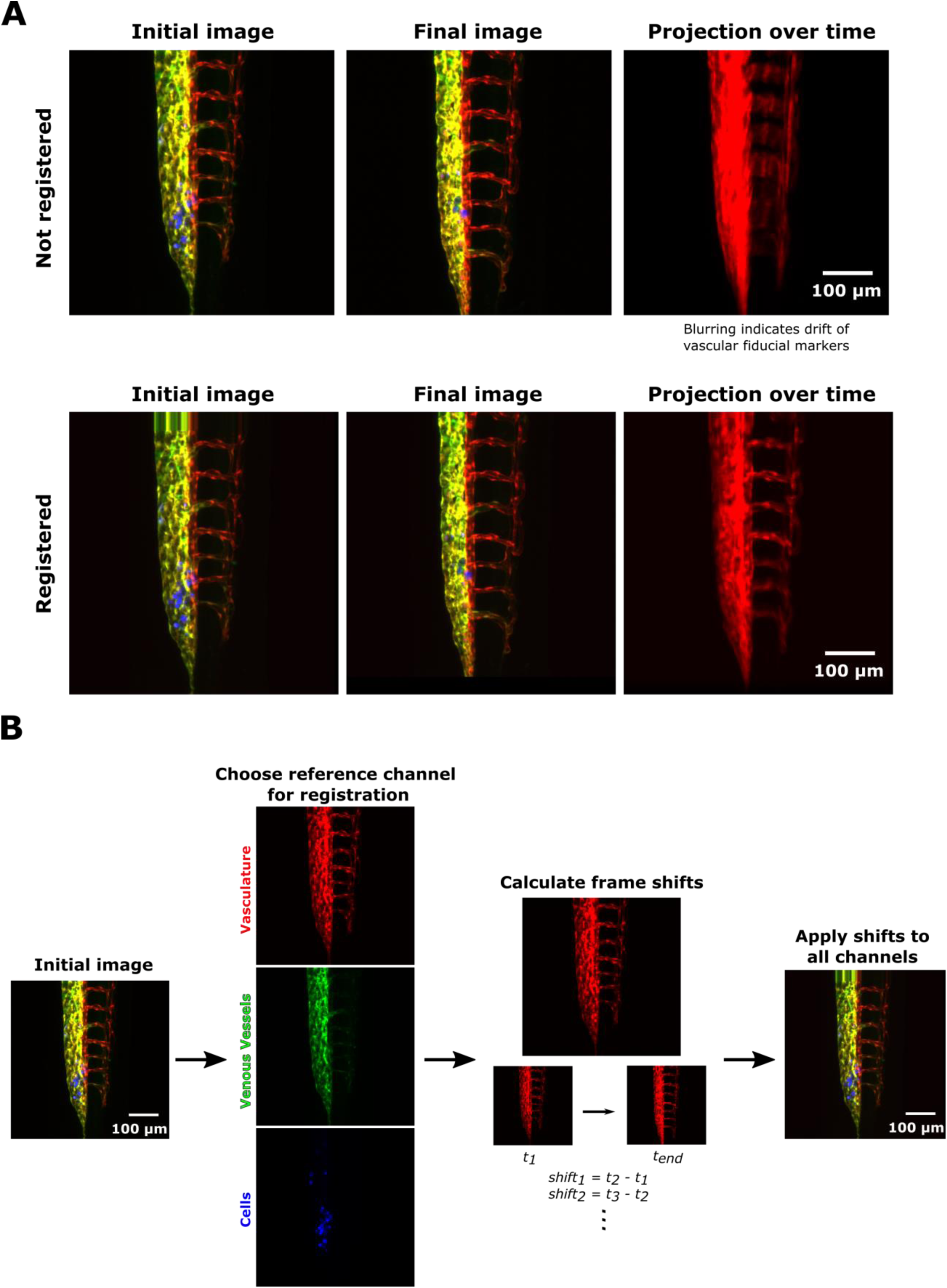
Related to Figure 3. Image registration to correct for zebrafish growth and drift during cell speed analysis. (A) Representative images of the zebrafish tail, with images taken from time-lapse imaging of cell movement. The initial (t=0 h) and final (t=12 h) frames are shown for the raw images (top) and registered images (bottom). An average intensity projection of the vasculature (shown in red) over time illustrates drift in the non-registered image. (B) Schematic of image registration process. A reference channel containing fiducial markers is chosen to register images over time. For each time interval, frame shifts are calculated. These shifts are then applied to all channels of the image stacks to obtain a registered image. In this figure, all images are average intensity projections of confocal z stacks. Registration was performed in 3D on z stacks prior to analysis. See also Figure 3.

**Figure S5.**
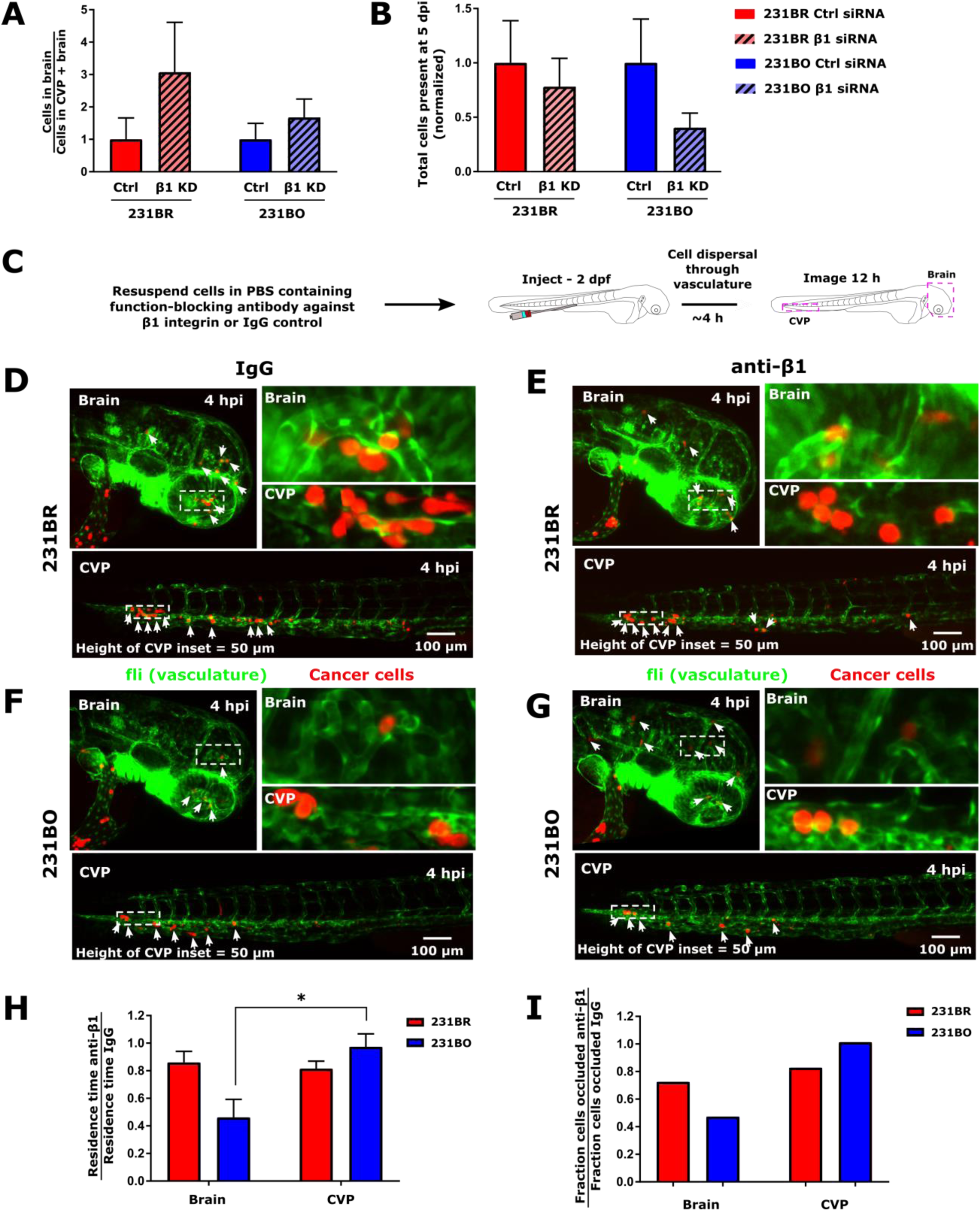
Related to Figure 5. Effect of β1 integrin knockdown and function blocking on cell survival and organ homing. (A) Ratio of cells homing to the brain to cells homing to cells present in the brain and CVP at 5 dpi, normalized against the average value for Ctrl cells for both MDA-MB-231 brain-tropic (231BR) and bone marrow-tropic (231BO) lines (mean ± SEM). Values were not significantly different by two-way ANOVA with Sidak’s multiple comparisons post-test. (B) Total cells present in the zebrafish brain and caudal vein plexus (CVP) at 5 days post-injection (dpi) upon knockdown of β1 integrin (β1 KD) or transfection with non-targeting control siRNA (Ctrl). Values were normalized to those in larvae injected with cells transfected with non-targeting control siRNA (mean ± SEM). Values were not significantly different by two-way ANOVA with Sidak’s multiple comparisons post-test within cell types. N=5 larvae for 231BR Ctrl cells, N=5 larvae for 231BR siβ1 cells, N=5 larvae for 231BO Ctrl cells, and N=5 larvae for 231BO siβ1 cells. (C) Schematic of β1 integrin function blocking experiment. MDA-MB-231 brain-tropic or bone marrow-tropic cells were incubated with a function-blocking antibody targeting β1 integrin or a non-targeting control IgG antibody. Cells were injected to the circulation of d days post-fertilization (dpf) Tg(fli:EGFP) zebrafish, and the brain and caudal vein plexus (CVP) were imaged over the first ~12 h following cell dispersal through the vasculature. Representative images of MDA-MB-231 brain-tropic (231BR) cells treated with (D) IgG control or (E) function blocking antibody against β1 integrin and MDA-MB-231 bone marrow-tropic (231BO) cells treated with (F) IgG control or (G) function blocking antibody against β1 integrin approximately 4 h after injection to 2 dpf Tg(fli:EGFP) zebrafish. Inset locations are indicated on the top left and bottom panels. Cells are displayed in red, and the vasculature is displayed in green. Arrows indicate cell positions. For scale, the height of the caudal vein plexus (CVP) inset = 50 μm. (H) Residence times (mean ± SEM) of brain-tropic 231BR and bone marrow-tropic 231BO cells treated with β1 integrin function blocking antibody in the brain and CVP, normalized by the average residence time of the same cell type treated with an IgG isotype control antibody in the same tissue. *, p<0.05 by Sidak’s multiple comparisons test between the same cell type following two-way ANOVA. (I) Fractional change in cell occlusion in each organ and for each cell type upon inhibition of β1 integrin. Fraction of cells present in each tissue after 12 h of tracking to total number of cells tracked were normalized to the fraction of the same cell type occluded upon treatment with IgG isotype control antibody. Residence times and occlusion fractions were calculated from: 231BR IgG – 5 larvae, 207 cells in head, 159 cells in CVP; 231BR anti-β1 – 4 larvae, 181 cells in head, 296 cells in CVP; 231BO IgG – 5 larvae, 78 cells in head, 139 cells in CVP; anti-β1 – 3 larvae, 26 cells in head, 77 cells in CVP. See also Figure 5.

**Supplementary Video 1.**
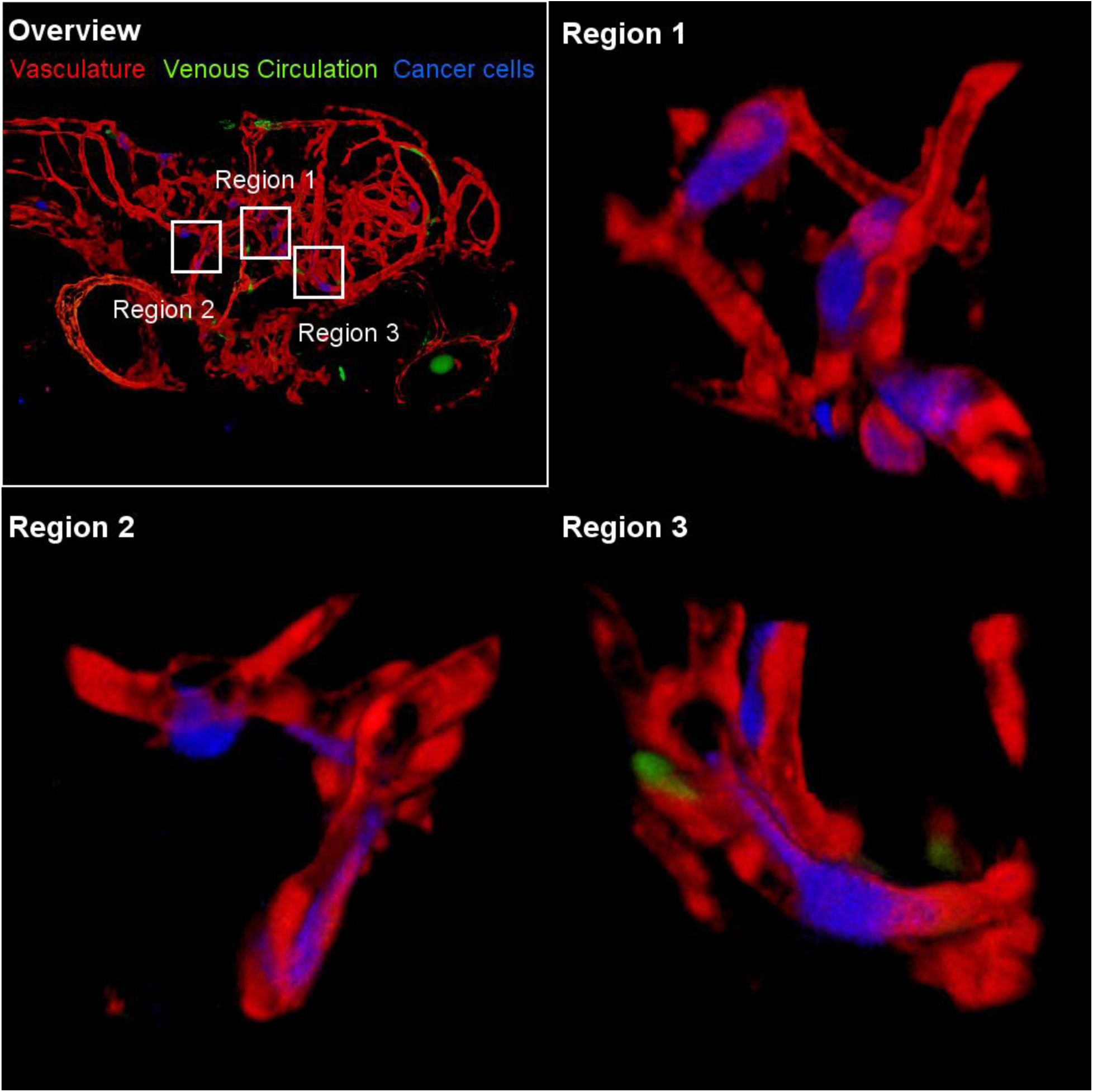
Related to Figure 1. Early colonization of the zebrafish brain by human breast cancer cells. Human brain-tropic MDA-MB-231 breast cancer cells (231BR) were injected into the circulation of 2 days post-fertilization (dpf) Tg(flk:mCherry/MRC1a:EGFP) zebrafish. The brain was imaged 5 days post-injection (dpi). 3D reconstructions from confocal z stacks of the zebrafish head and three regions brain are shown. Extravasated and intravascular cells are indicated. See also Figure 1.

**Supplementary Video 2.**
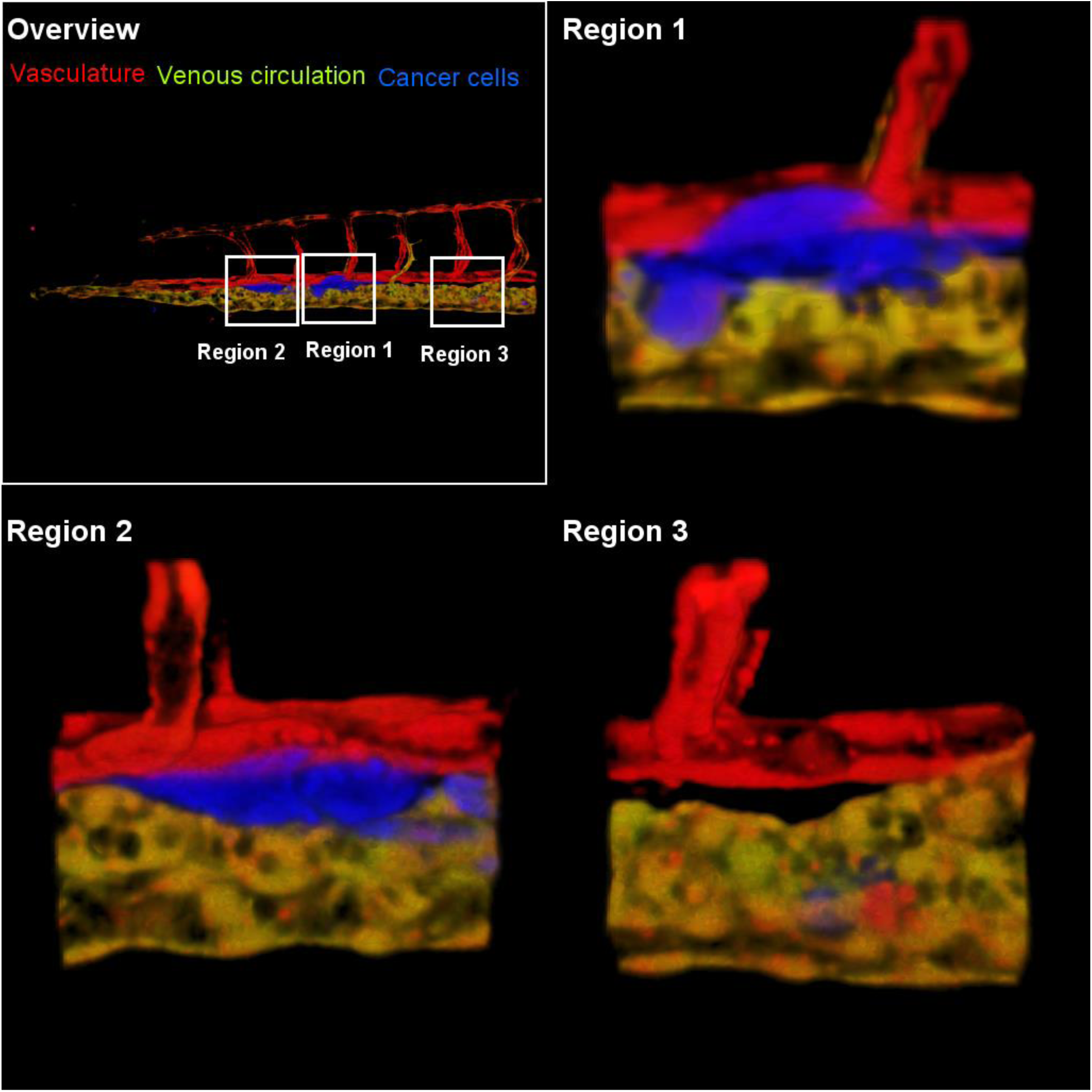
Related to Figure 1. Early colonization of the zebrafish caudal vein plexus by human breast cancer cells. Human bone marrow-tropic MDA-MB-231 breast cancer cells (231BO) were injected into the circulation of 2 days post-fertilization (dpf) Tg(flk:mCherry/MRC1a:EGFP) zebrafish. The brain was imaged 5 days post-injection (dpi). 3D reconstructions from confocal z stacks of the zebrafish caudal vein plexus (CVP) shown, with three regions of interest highlighted and shown in detail. Extravasated and intravascular cells are indicated. See also Figure 1.

**Supplementary Video 3.**
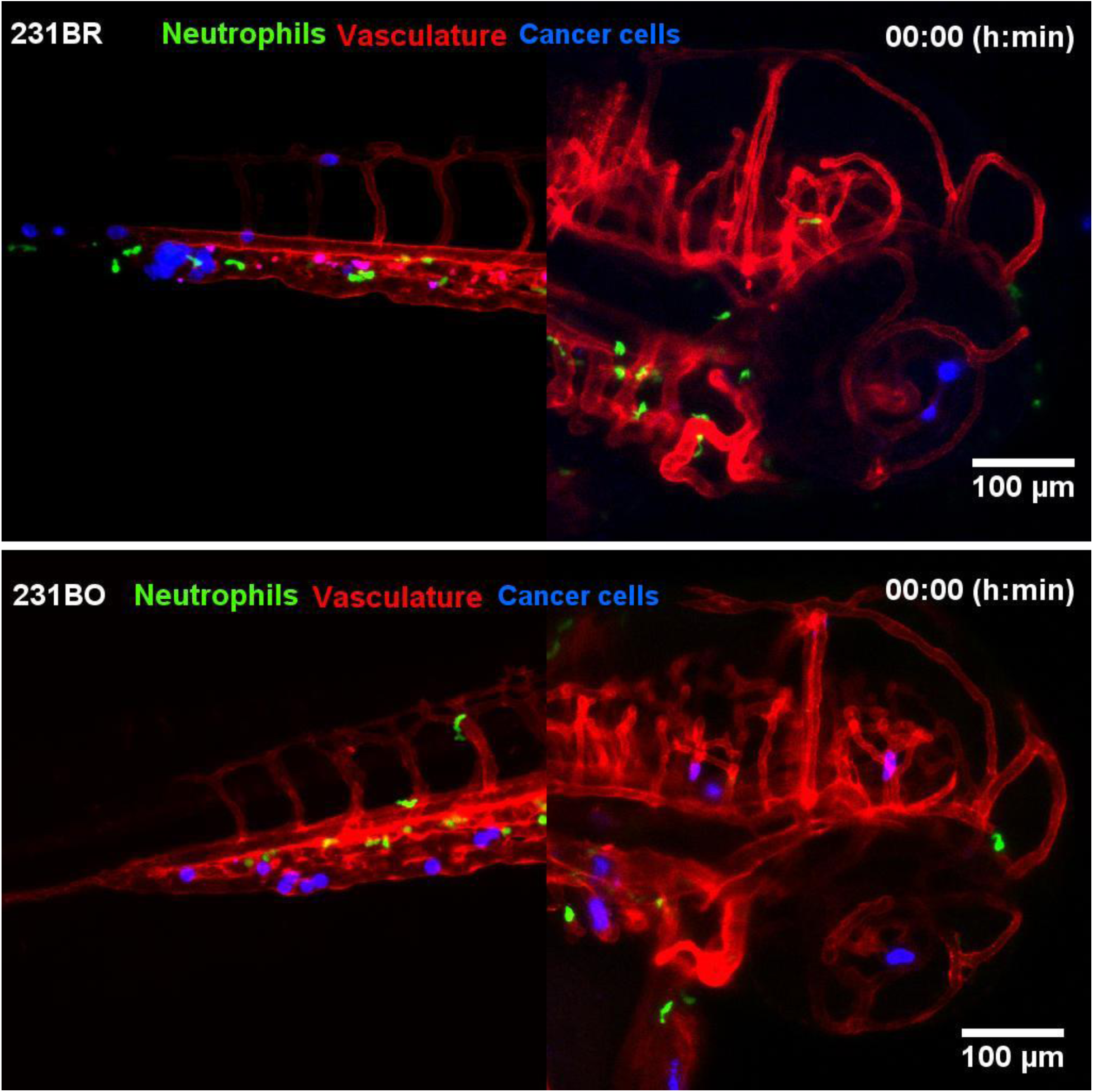
Related to Figure 2. Neutrophils do not make prolonged contact with disseminated cancer cells. 231BR and 231BO cells (displayed in blue) are shown moving through the vasculature of 2–3 dpf Tg(mpx:EGFP/flk:mCherry) zebrafish following injection to the circulation. The vasculature is displayed in red, and zebrafish neutrophils are displayed in green. Images were acquired every 10 min. Video is average intensity projection of confocal z stack. See also Figure 2.

**Supplementary Video 4.**
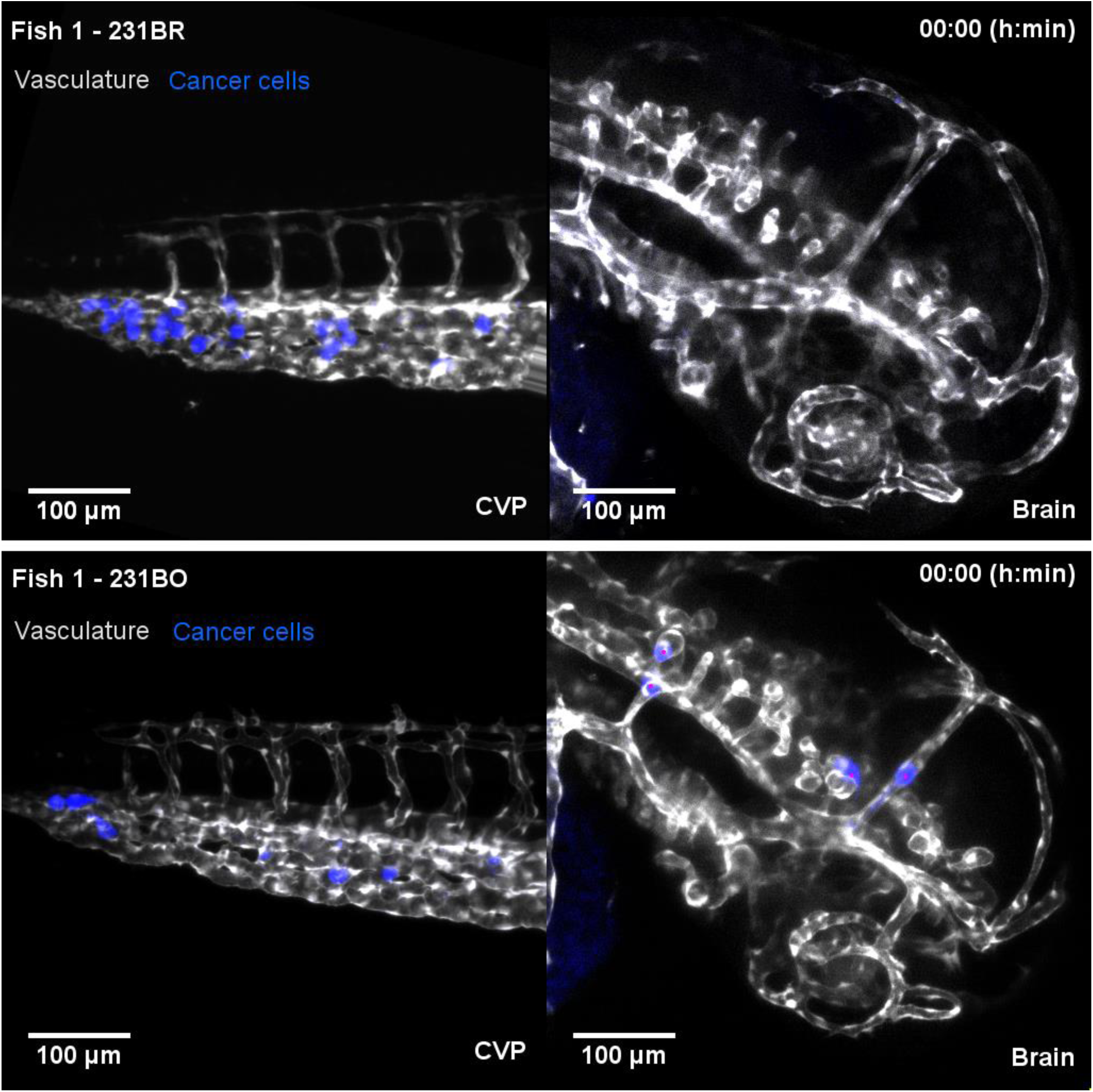
Related to Figure 3. Intravital cell tracking of MDA-MB-231 organ-tropic cells in the zebrafish brain and caudal vein plexus (CVP). MDA-MB-231 brain-tropic (231BR) or bone marrow-tropic (231BO) cells (displayed in blue) are shown moving through the vasculature of 2–3 days post-fertilization (dpf) Tg(flk:mCherry/MRC1a:EGFP) zebrafish following injection to the circulation. Zebrafish vasculature and venous circulation are shown in grayscale to facilitate visualization of cancer cell trafficking. Cell trajectories are displayed as colored lines. Images were acquired every 10 min. Video is average intensity projection of confocal z stack, and cells were tracked in 3 dimensions. See also Figure 3.

**Supplementary Video 5.**
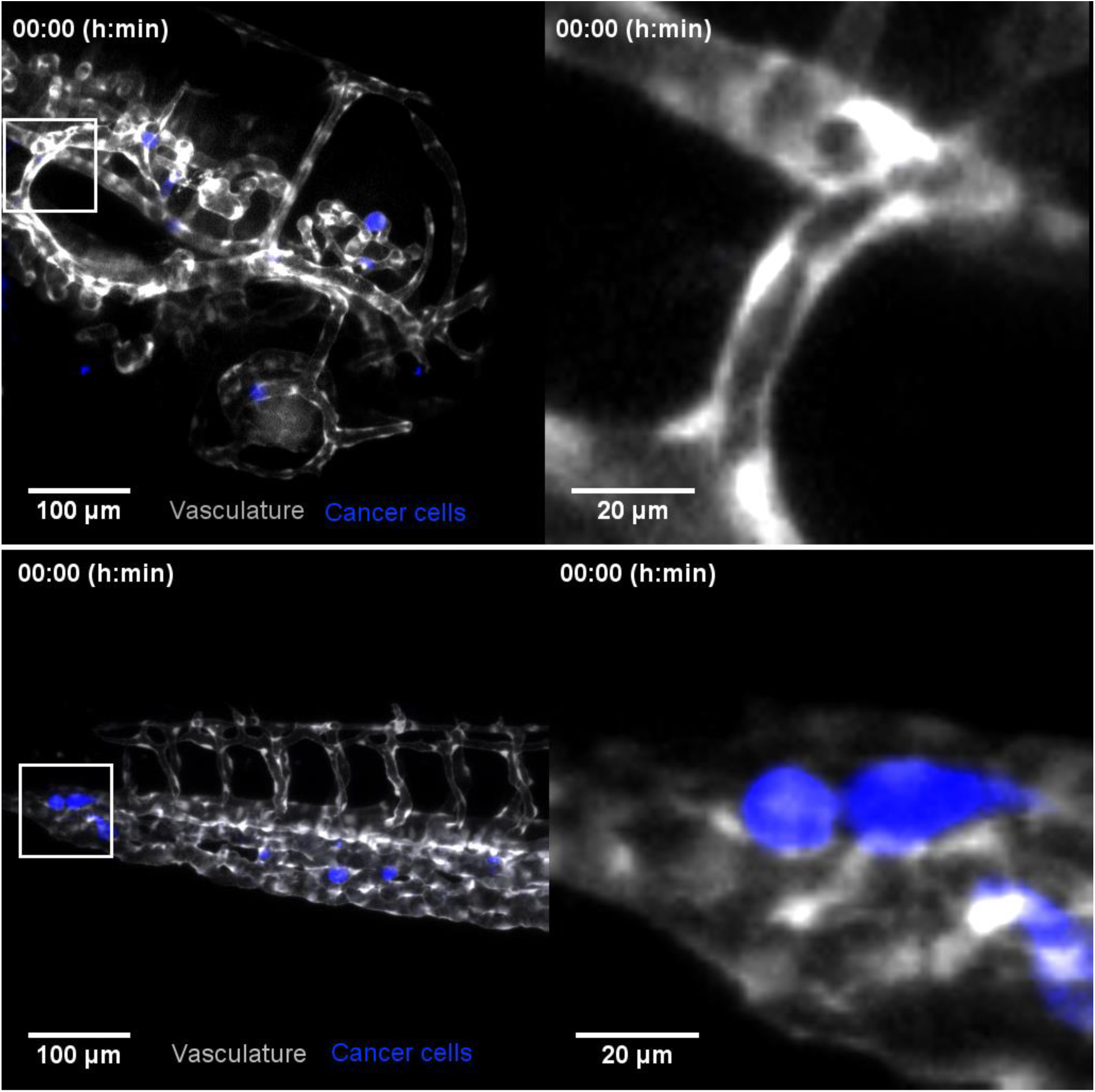
Related to Figure 4. Tumor cell extravasation in the zebrafish brain and CVP. Top panels - MDA-MB-231 brain-tropic (231BR) cell extravasation in the brain of 2–3 days post-fertilization (dpf) Tg(flk:mCherry/MRC1a:EGFP) zebrafish following injection to the circulation. Bottom panels - MDA-MB-231 bone marrow-tropic (231BO) cell extravasation in the caudal vein plexus (CVP) of 2–3 days post-fertilization (dpf) Tg(flk:mCherry/MRC1a:EGFP) zebrafish following injection to the circulation. Insets shows detail of extravasating cell. Images were acquired every 10 min on the day of cell injection. Images are average intensity projections of confocal z stack. See also Figure 4.

## REFERENCES

Azevedo, A.S., Follain, G., Patthabhiraman, S., Harlepp, S., and Goetz, J.G. (2015). Metastasis of circulating tumor cells: favorable soil or suitable biomechanics, or both? Cell Adh Migr 9, 345–356.

Balzer, E.M., Tong, Z., Paul, C.D., Hung, W.C., Stroka, K.M., Boggs, A.E., Martin, S.S., and Konstantopoulos, K. (2012). Physical confinement alters tumor cell adhesion and migration phenotypes. FASEB journal: official publication of the Federation of American Societies for Experimental Biology 26, 4045–4056.

Barnes, J.M., Przybyla, L., and Weaver, V.M. (2017). Tissue mechanics regulate brain development, homeostasis and disease. J Cell Sci 130, 71–82.

Bissell, M.J., and Hines, W.C. (2011). Why don’t we get more cancer? A proposed role of the microenvironment in restraining cancer progression. Nat Med 17, 320–329.

Blehm, B.H., Devine, A., Staunton, J.R., and Tanner, K. (2016). In vivo tissue has non-linear rheological behavior distinct from 3D biomimetic hydrogels, as determined by AMOTIV microscopy. Biomaterials 83, 66–78.

Chen, M.B., Lamar, J.M., Li, R., Hynes, R.O., and Kamm, R.D. (2016). Elucidation of the Roles of Tumor Integrin beta1 in the Extravasation Stage of the Metastasis Cascade. Cancer Res 76, 2513–2524.

Condeelis, J., and Segall, J.E. (2003). Intravital imaging of cell movement in tumours. Nat Rev Cancer 3, 921–930.

Cox, J., Hein, M.Y., Luber, C.A., Paron, I., Nagaraj, N., and Mann, M. (2014). Accurate proteome-wide label-free quantification by delayed normalization and maximal peptide ratio extraction, termed MaxLFQ. Molecular & cellular proteomics: MCP 13, 2513–2526.

Cox, J., and Mann, M. (2008). MaxQuant enables high peptide identification rates, individualized p.p.b.-range mass accuracies and proteome-wide protein quantification. Nature biotechnology 26, 1367–1372.

Doerschuk, C.M., Beyers, N., Coxson, H.O., Wiggs, B., and Hogg, J.C. (1993). Comparison of neutrophil and capillary diameters and their relation to neutrophil sequestration in the lung. Journal of applied physiology (Bethesda, Md: 1985) 74, 3040–3045.

Ewald, A.J., Werb, Z., and Egeblad, M. (2011). Dynamic, long-term in vivo imaging of tumor-stroma interactions in mouse models of breast cancer using spinning-disk confocal microscopy. Cold Spring Harb Protoc 2011, pdb top97.

Fidler, I.J. (2003). The pathogenesis of cancer metastasis: the ‘seed and soil’ hypothesis revisited. Nat Rev Cancer 3, 453–458.

Fischer, M., and Berg-Sørensen, K. (2007). Calibration of trapping force and response function of optical tweezers in viscoelastic media. Journal of Optics A: Pure and Applied Optics 9, S239.

Fleming, A., Diekmann, H., and Goldsmith, P. (2013). Functional characterisation of the maturation of the blood-brain barrier in larval zebrafish. PloS one 8, e77548.

Follain, G., Osmani, N., Azevedo, A.S., Allio, G., Mercier, L., Karreman, M.A., Solecki, G., Garcia Leon, M.J., Lefebvre, O., Fekonja, N., et al. (2018). Hemodynamic Forces Tune the Arrest, Adhesion, and Extravasation of Circulating Tumor Cells. Developmental cell 45, 33–52.e12.

Fonck, E., Feigl, G.G., Fasel, J., Sage, D., Unser, M., Rufenacht, D.A., and Stergiopulos, N. (2009). Effect of aging on elastin functionality in human cerebral arteries. Stroke 40, 2552–2556.

Friedl, P., and Alexander, S. (2011). Cancer invasion and the microenvironment: plasticity and reciprocity. Cell 147, 992–1009.

Goessling, W., and North, T.E. (2011). Hematopoietic stem cell development: using the zebrafish to identify the signaling networks and physical forces regulating hematopoiesis. Methods in cell biology 105, 117–136.

Hirata, E., Girotti, M.R., Viros, A., Hooper, S., Spencer-Dene, B., Matsuda, M., Larkin, J., Marais, R., and Sahai, E. (2015). Intravital imaging reveals how BRAF inhibition generates drug-tolerant microenvironments with high integrin beta1/FAK signaling. Cancer Cell 27, 574–588.

Hoshino, A., Costa-Silva, B., Shen, T.L., Rodrigues, G., Hashimoto, A., Tesic Mark, M., Molina, H., Kohsaka, S., Di Giannatale, A., Ceder, S., et al. (2015). Tumour exosome integrins determine organotropic metastasis. Nature 527, 329–335.

Jung, H.M., Castranova, D., Swift, M.R., Pham, V.N., Venero Galanternik, M., Isogai, S., Butler, M.G., Mulligan, T.S., and Weinstein, B.M. (2017). Development of the larval lymphatic system in zebrafish. Development 144, 2070–2081.

Kennecke, H., Yerushalmi, R., Woods, R., Cheang, M.C., Voduc, D., Speers, C.H., Nielsen, T.O., and Gelmon, K. (2010). Metastatic behavior of breast cancer subtypes. J Clin Oncol 28, 3271–3277.

Kienast, Y., von Baumgarten, L., Fuhrmann, M., Klinkert, W.E., Goldbrunner, R., Herms, J., and Winkler, F. (2010). Real-time imaging reveals the single steps of brain metastasis formation. Nat Med 16, 116–122.

Kim, J., Staunton, J.R., and Tanner, K. (2016). Independent Control of Topography for 3D Patterning of the ECM Microenvironment. Advanced materials (Deerfield Beach, Fla) 28, 132–137.

Kim, J., and Tanner, K. (2015). Recapitulating the Tumor Ecosystem Along the Metastatic Cascade Using 3D Culture Models. Front Oncol 5, 170.

Kim, S.H., Redvers, R.P., Chi, L.H., Ling, X., Lucke, A.J., Reid, R.C., Fairlie, D.P., Baptista Moreno Martin, A.C., Anderson, R.L., Denoyer, D., et al. (2018). Identification of brain metastasis genes and therapeutic evaluation of histone deacetylase inhibitors in a clinically relevant model of breast cancer brain metastasis. Dis Model Mech.

Kniazeva, E., Weidling, J.W., Singh, R., Botvinick, E.L., Digman, M.A., Gratton, E., and Putnam, A.J. (2012). Quantification of local matrix deformations and mechanical properties during capillary morphogenesis in 3D. Integr Biol (Camb) 4, 431–439.

Kotlarchyk, M.A., Shreim, S.G., Alvarez-Elizondo, M.B., Estrada, L.C., Singh, R., Valdevit, L., Kniazeva, E., Gratton, E., Putnam, A.J., and Botvinick, E.L. (2011). Concentration independent modulation of local micromechanics in a fibrin gel. PloS one 6, e20201.

Kumar, S., and Weaver, V.M. (2009). Mechanics, malignancy, and metastasis: the force journey of a tumor cell. Cancer Metastasis Rev 28, 113–127.

Kusuma, N., Denoyer, D., Eble, J.A., Redvers, R.P., Parker, B.S., Pelzer, R., Anderson, R.L., and Pouliot, N. (2012). Integrin-dependent response to laminin-511 regulates breast tumor cell invasion and metastasis. International journal of cancer 130, 555–566.

Kusumbe, A.P., Ramasamy, S.K., and Adams, R.H. (2014). Coupling of angiogenesis and osteogenesis by a specific vessel subtype in bone. Nature 507, 323–328.

Lawson, N.D., and Weinstein, B.M. (2002). In vivo imaging of embryonic vascular development using transgenic zebrafish. Developmental biology 248, 307–318.

Ma, D., Zhang, J., Lin, H.F., Italiano, J., and Handin, R.I. (2011). The identification and characterization of zebrafish hematopoietic stem cells. Blood 118, 289–297.

Mathieu, E., Paul, C.D., Stahl, R., Vanmeerbeeck, G., Reumers, V., Liu, C., Konstantopoulos, K., and Lagae, L. (2016). Time-lapse lens-free imaging of cell migration in diverse physical microenvironments. Lab on a chip 16, 3304–3316.

Mathura, K.R., Vollebregt, K.C., Boer, K., De Graaff, J.C., Ubbink, D.T., and Ince, C. (2001). Comparison of OPS imaging and conventional capillary microscopy to study the human microcirculation. Journal of applied physiology (Bethesda, Md: 1985) 91, 74–78.

Meeker, N.D., and Trede, N.S. (2008). Immunology and zebrafish: spawning new models of human disease. Dev Comp Immunol 32, 745–757.

Murayama, E., Kissa, K., Zapata, A., Mordelet, E., Briolat, V., Lin, H.F., Handin, R.I., and Herbomel, P. (2006). Tracing hematopoietic precursor migration to successive hematopoietic organs during zebrafish development. Immunity 25, 963–975.

Neuman, K.C., and Block, S.M. (2004). Optical trapping. Rev Sci Instrum 75, 2787–2809.

Nguyen-Chi, M., Laplace-Builhe, B., Travnickova, J., Luz-Crawford, P., Tejedor, G., Phan, Q.T., Duroux-Richard, I., Levraud, J.-P., Kissa, K., Lutfalla, G., et al. (2015). Identification of polarized macrophage subsets in zebrafish. eLife 4, e07288.

Nguyen, D.X., Bos, P.D., and Massague, J. (2009). Metastasis: from dissemination to organ-specific colonization. Nat Rev Cancer 9, 274–284.

Nombela-Arrieta, C., Pivarnik, G., Winkel, B., Canty, K.J., Harley, B., Mahoney, J.E., Park, S.Y., Lu, J., Protopopov, A., and Silberstein, L.E. (2013). Quantitative imaging of haematopoietic stem and progenitor cell localization and hypoxic status in the bone marrow microenvironment. Nature cell biology 15, 533–543.

Obenauf, A.C., and Massague, J. (2015). Surviving at a distance: organ specific metastasis. Trends Cancer 1, 76–91.

Oosterhof, N., Boddeke, E., and van Ham, T.J. (2015). Immune cell dynamics in the CNS: Learning from the zebrafish. Glia 63, 719–735.

Oudin, M.J., and Weaver, V.M. (2016). Physical and Chemical Gradients in the Tumor Microenvironment Regulate Tumor Cell Invasion, Migration, and Metastasis. Cold Spring Harb Symp Quant Biol 81, 189–205.

Paget, S. (1989). The distribution of secondary growths in cancer of the breast. 1889. Cancer Metastasis Rev 8, 98–101.

Parslow, A., Cardona, A., and Bryson-Richardson, R.J. (2014). Sample drift correction following 4D confocal time-lapse imaging. Journal of visualized experiments: JoVE.

Patsialou, A., Wyckoff, J., Wang, Y., Goswami, S., Stanley, E.R., and Condeelis, J.S. (2009). Invasion of human breast cancer cells in vivo requires both paracrine and autocrine loops involving the colony-stimulating factor-1 receptor. Cancer Res 69, 9498–9506.

Paul, C.D., Mistriotis, P., and Konstantopoulos, K. (2017). Cancer cell motility: lessons from migration in confined spaces. Nature reviews Cancer 17, 131–140.

Potter, R.F., and Groom, A.C. (1983). Capillary diameter and geometry in cardiac and skeletal muscle studied by means of corrosion casts. Microvascular research 25, 68–84.

Puspoki, Z., Storath, M., Sage, D., and Unser, M. (2016). Transforms and Operators for Directional Bioimage Analysis: A Survey. Advances in anatomy, embryology, and cell biology 219, 69–93.

Ramasamy, S.K., Kusumbe, A.P., Schiller, M., Zeuschner, D., Bixel, M.G., Milia, C., Gamrekelashvili, J., Limbourg, A., Medvinsky, A., Santoro, M.M., et al. (2016). Blood flow controls bone vascular function and osteogenesis. Nature communications 7, 13601.

Renshaw, S.A., and Trede, N.S. (2012). A model 450 million years in the making: zebrafish and vertebrate immunity. Dis Model Mech 5, 38–47.

Sacco, A., Roccaro, A.M., Ma, D., Shi, J., Mishima, Y., Moschetta, M., Chiarini, M., Munshi, N., Handin, R.I., and Ghobrial, I.M. (2016). Cancer Cell Dissemination and Homing to the Bone Marrow in a Zebrafish Model. Cancer Res 76, 463–471.

Schwartz, M.A. (2010). Integrins and extracellular matrix in mechanotransduction. Cold Spring Harb Perspect Biol 2, a005066.

Staunton, J.R., Blehm, B., Devine, A., and Tanner, K. (2017). In situ calibration of position detection in an optical trap for active microrheology in viscous materials. Opt Express 25, 1746–1761.

Staunton, J.R., Vieira, W., Fung, K.L., Lake, R., Devine, A., and Tanner, K. (2016a). Mechanical properties of the tumor stromal microenvironment probed in vitro and ex vivo by in situ-calibrated optical trap-based active microrheology. Cell Mol Bioeng 9, 398–417.

Staunton, J.R., Vieira, W., Fung, K.L., Lake, R., Devine, A., and Tanner, K. (2016b). Mechanical properties of the tumor stromal microenvironment probed in vitro and ex vivo by in situ-calibrated optical trap-based active microrheology. Cellular and molecular bioengineering 9, 398–417.

Steeg, P.S. (2016). Targeting metastasis. Nat Rev Cancer 16, 201–218.

Steeg, P.S., Camphausen, K.A., and Smith, Q.R. (2011). Brain metastases as preventive and therapeutic targets. Nat Rev Cancer 11, 352–363.

Stoletov, K., Kato, H., Zardouzian, E., Kelber, J., Yang, J., Shattil, S., and Klemke, R. (2010). Visualizing extravasation dynamics of metastatic tumor cells. J Cell Sci 123, 2332–2341.

Stoletov, K., Montel, V., Lester, R.D., Gonias, S.L., and Klemke, R. (2007). High-resolution imaging of the dynamic tumor cell vascular interface in transparent zebrafish. Proc Natl Acad Sci U S A 104, 17406–17411.

Stroka, Kimberly M., Jiang, H., Chen, S.-H., Tong, Z., Wirtz, D., Sun, Sean X., and Konstantopoulos, K. (2014). Water Permeation Drives Tumor Cell Migration in Confined Microenvironments. Cell 157, 611–623.

Tanner, K., and Gottesman, M.M. (2015). Beyond 3D culture models of cancer. Sci Transl Med 7, 283ps289.

Tanner, K., Mori, H., Mroue, R., Bruni-Cardoso, A., and Bissell, M.J. (2012). Coherent angular motion in the establishment of multicellular architecture of glandular tissues. Proc Natl Acad Sci U S A 109, 1973–1978.

Tinevez, J.Y., Perry, N., Schindelin, J., Hoopes, G.M., Reynolds, G.D., Laplantine, E., Bednarek, S.Y., Shorte, S.L., and Eliceiri, K.W. (2017). TrackMate: An open and extensible platform for single-particle tracking. Methods (San Diego, Calif) 115, 80–90.

Tufto, I., and Rofstad, E.K. (1999). Interstitial fluid pressure and capillary diameter distribution in human melanoma xenografts. Microvascular research 58, 205–214.

Tyanova, S., Temu, T., Sinitcyn, P., Carlson, A., Hein, M.Y., Geiger, T., Mann, M., and Cox, J. (2016). The Perseus computational platform for comprehensive analysis of (prote)omics data. Nature methods 13, 731–740.

Watkins, S.C., Maniar, S., Mosher, M., Roman, B.L., Tsang, M., and St Croix, C.M. (2012). High resolution imaging of vascular function in zebrafish. PloS one 7, e44018.

Weaver, V.M., Petersen, O.W., Wang, F., Larabell, C.A., Briand, P., Damsky, C., and Bissell, M.J. (1997). Reversion of the malignant phenotype of human breast cells in three-dimensional culture and in vivo by integrin blocking antibodies. J Cell Biol 137, 231–245.

White, R., Rose, K., and Zon, L. (2013). Zebrafish cancer: the state of the art and the path forward. Nat Rev Cancer 13, 624–636.

Wisniewski, J.R., Zougman, A., Nagaraj, N., and Mann, M. (2009). Universal sample preparation method for proteome analysis. Nature methods 6, 359–362.

Wyckoff, J.B., Wang, Y., Lin, E.Y., Li, J.F., Goswami, S., Stanley, E.R., Segall, J.E., Pollard, J.W., and Condeelis, J. (2007). Direct visualization of macrophage-assisted tumor cell intravasation in mammary tumors. Cancer Res 67, 2649–2656.

Yoneda, T., Williams, P.J., Hiraga, T., Niewolna, M., and Nishimura, R. (2001). A bone-seeking clone exhibits different biological properties from the MDA-MB-231 parental human breast cancer cells and a brain-seeking clone in vivo and in vitro. J Bone Miner Res 16, 1486–1495.

Zhang, S., Huang, W.C., Zhang, L., Zhang, C., Lowery, F.J., Ding, Z., Guo, H., Wang, H., Huang, S., Sahin, A.A., et al. (2013). SRC family kinases as novel therapeutic targets to treat breast cancer brain metastases. Cancer Res 73, 5764–5774.

